# The retrosplenial cortex combines internal and external cues to encode head velocity during navigation

**DOI:** 10.1101/2021.01.22.427789

**Authors:** Sepiedeh Keshavarzi, Edward F. Bracey, Richard A. Faville, Dario Campagner, Adam L. Tyson, Stephen C. Lenzi, Tiago Branco, Troy W. Margrie

## Abstract

The extent to which we successfully navigate the environment depends on our ability to continuously track our heading direction and speed. Angular head velocity (AHV) cells, which encode the speed and direction of head turns during navigation, are fundamental to this process, yet the mechanisms that determine their function remain unknown. By performing chronic single-unit recordings in the retrosplenial cortex (RSP) of the mouse and tracking the activity of individual AHV neurons between freely moving and head-restrained conditions, we find that vestibular inputs dominate AHV signalling. In addition, we discover that self-generated optic flow input onto these neurons increases the gain and signal-to-noise ratio of angular velocity coding during free exploration. Psychophysical experiments and neural decoding further reveal that vestibular-visual integration increases the perceptual accuracy of egocentric angular velocity and the fidelity of its representation by RSP ensembles. We propose that while AHV coding is dependent on vestibular input, it also uses vision to maximise navigation accuracy in nocturnal and diurnal environments.

## Introduction

Whether foraging for food or taking refuge from danger, the survival of most animals depends on their knowledge of current location and heading direction. Such cognitive capacity requires not only an accurate estimation of the momentary heading direction, but also the ability to update it with subsequent head turns as the animal moves through space (1–3). Head direction cells, identified in various animals ranging from insects to primates, are tuned to the azimuthal direction of an animal’s head in its environment (4–6), and are thought to provide an internal representation of the sense of direction (7, 8). It is widely accepted that the updating of this heading direction signal arises from angular velocity integration (1–3, 9, 10), with theoretical models of head direction computation requiring the existence of neurons that encode angular head velocity (AHV, (11, 12)) a) and neurons with conjunctive head direction and AHV tuning (13–16).

In theory, AHV information can be obtained from multiple internal and external sources, including vestibular signals, efference copy of motor commands, proprioceptive feedback from eye and neck muscles, and optic flow; yet, whether and how these motion cues contribute to AHV coding during navigation remains unknown. On one hand, perceptual experiments in humans and non-human primates suggest that the combination of vestibular and visual stimuli improves estimates of heading during passive translation (17–21). On the other hand, for instance in non-cortical areas, vestibular signals are suggested to be either completely inhibited or only weakly active during self-generated movements (12, 22–24). In spite of this, successful path integration and head direction tuning in the thalamus appears to require an intact vestibular system (25–27).

A major obstacle impeding progress in understanding AHV coding and other egocentric elements of spatial navigation, is the inability to precisely disentangle the contribution of these various sensory cues in freely moving animals. Conversely, while head-fixed experiments allow quantitative control of sensory stimuli, the direct relevance of signals recorded in such preparations to natural navigation is unclear. Here, we have identified cortical AHV cells during free exploration and, for the first time, determined their response properties to isolated and conjunctive sensory stimuli. We have focused on AHV cells in the RSP since it is a multisensory cortical region that forms part of the head direction network and encodes various spatial and movement-related information (28–38). Notably, studies on both rodents and humans suggest that the RSP is critically involved in spatial orientation and self motion-guided navigation (39–43), and recent work suggests that it can provide head-motion information to the visual cortex (44). A growing number of studies in recent years has advanced our knowledge of allocentric head direction representation in the RSP and the potential role of this region in processing landmark information (32, 45–48). However, despite increasing evidence for egocentric computations in the RSP (34, 35, 38), the extent to which self-motion information such as AHV is represented in this region and the underlying mechanisms remains unexplored.

## Results

Using chronically implanted high-density silicone probes (Neuropixels and Neuronexus), RSP neurons (n = 359 single units in 5 mice, right hemisphere) were first recorded from mice while they freely explored a circular arena under both light and dark conditions (Fig. 1A, Fig. S1A). Head and body movements were tracked in 2D (horizontal plane of motion, Fig. 1A) and, based on their tuning function, neurons were classified as encoding one, or a combination, of three navigational properties: heading direction, AHV, and linear locomotion speed (Fig. S1B, Fig. S2). Similar to previous reports (28, 29, 32), approximately 10 % of RSP neurons were identified as classic head direction cells (36/359, Fig. 1B, Fig. S2A, D). Strikingly, the majority (approximately 62 %) of recorded neurons were modulated by self-motion parameters including AHV (224/359, Fig. 1B, Fig. S2B, D-E), and – to a lesser degree – locomotion speed (182/359, Fig. 1B, Fig. S2C-D). In addition, many head direction cells also conjunctively encoded AHV (22/36, Fig. 1B). The spiking activity of most AHV cells was positively correlated with the speed of head turns (193/224, Fig. 1B). In 57 % of these cells (128/224), speed modulation was unidirectional, meaning that their firing rate was significantly correlated with the speed of either right or left head turns (Fig. S2B, middle), with no apparent preference for either direction at the population level (S2E). Additionally, a small fraction of cells (3/224, Fig. S2B, right) showed a positive speed correlation in one direction and a negative correlation in the other. Thus, AHV neurons in the RSP appear to represent both the speed and the direction of angular head movements. AHV cells were found across cortical layers in both dysgranular (105/224) and granular (119/224) divisions of the RSP (Fig. S1B), and consisted of both putative excitatory (121/224) and inhibitory (103/224) cell types (Fig. S3), suggesting that AHV representation in the RSP is widespread, distributed across regions, and more prevalent than previously reported (29, 49).

**Fig. 1.**
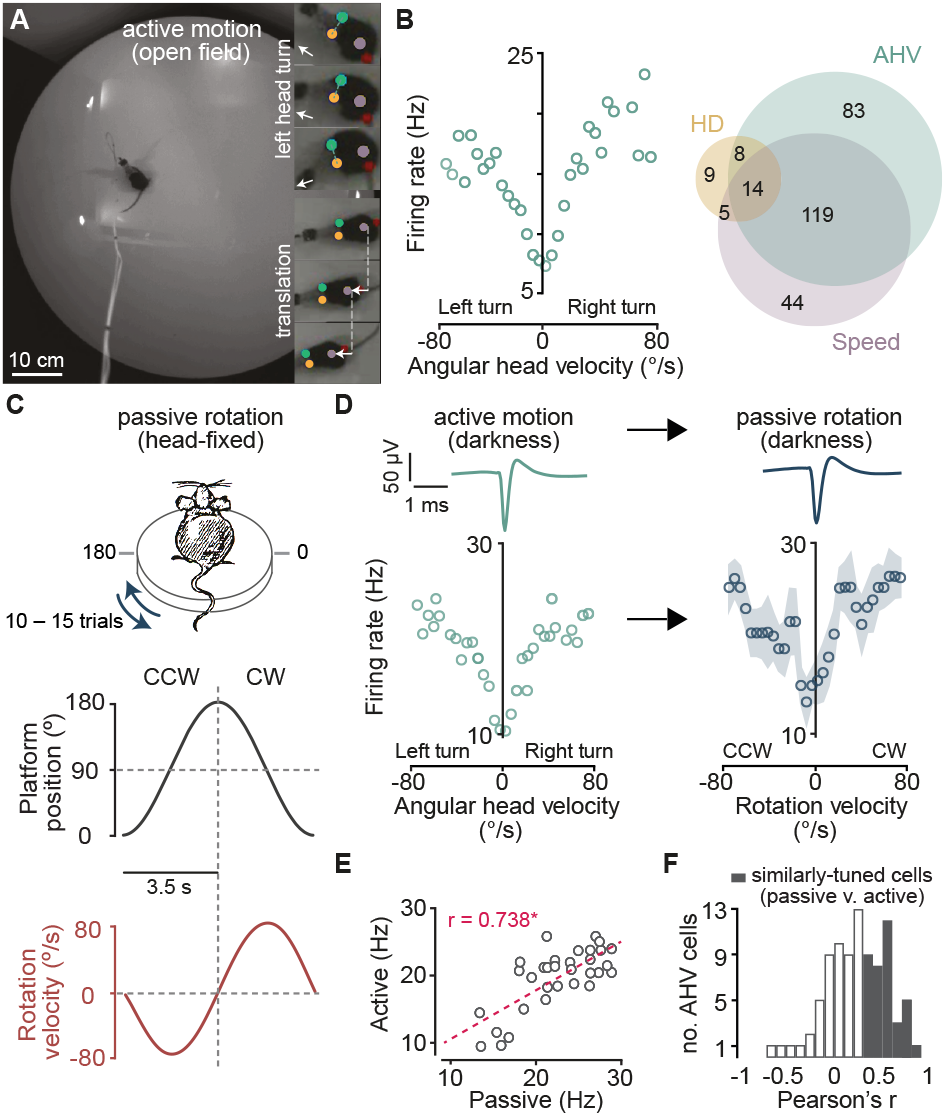
RSP neurons encode angular head velocity during both active exploration and passive motion. **(A)** A video frame showing the open-field arena used for exploration. Insets, video frames showing a left head turn (top three) and locomotion (bottom three) tracked using the position of ears and the body. Arrows indicate head direction (top three) and linear displacement (bottom three). **(B)** Left, tuning plot of a representative bidirectional AHV cell. Right, Venn diagram showing cells tuned to HD, AHV, and linear locomotion speed (n total = 359). **(C)** Top, schematic of the design for passive rotation experiments. Centre and bottom, position and velocity profile of an individual rotation stimulus. **(D)** Tuning plots of a bidirectional AHV cell recorded initially in the open-field (left), and subsequently during passive rotation (right) in darkness. Top, average trace of this unit’s spike waveform recorded under each condition. Circles and shaded area on the right show trial-averaged firing rates (12 trials) and SEM, respectively. **(E)** Firing rate at each velocity bin for the cell shown in D plotted for active versus passive conditions. * Pearson’s r > 95th percentile of the null distribution = similarly tuned. **(F)** Distribution of Pearson’s r of correlation between passive and active tuning curves for all tracked AHV neurons in the dark (n = 90).

To distinguish the contribution of various external and internal cues to AHV signalling, we tracked the same unit from the open field arena to a body- and head-restrained paradigm (Fig. S4) where we could parametrically vary vestibular and visual stimuli (Fig. 1C), and exclude any input from locomotion, neck-on-body proprioception, and voluntary head movements. To determine the contribution of vestibular signals, we first tested whether in the absence of visual input, AHV neurons recorded in the open field could encode the angular velocity of horizontal passive rotation. Out of 90 AHV neurons that were successfully tracked in head-fixed recordings, the spiking rate of 72 cells (80 %) was significantly correlated with the velocity of passive rotation in the dark. Interestingly, despite the initiation of complex head and body movements during free exploration as compared to head-fixed horizontal rotations, a substantial fraction of AHV neurons (48.8 ± 6.6 %, n = 5 mice, 12 recording sessions) had similar tuning to passive rotation (Fig. 1D-F), and this similarity was observed in all AHV tuning types (Fig. S5).

Considering the absence of efference copy and proprioception from voluntary head and neck movements and behavioural confounds such as postural signals (50), these findings indicate that, at least in the dark, AHV coding is primarily driven by vestibular input. In support of this, bilateral lesions of the horizontal and posterior semi-circular canals (Fig. S6A-B) significantly reduced rotation-evoked responses (Fig. S6C) and the magnitude of angular velocity correlations (Lesioned: median velocity Pearson’s r = 0.26, IQR = [0.12 – 0.42], n = 4 mice, 7 recordings, 313 cells; Control: median velocity Pearson’s r = 0.39, IQR = [0.2-0.6], n = 10 mice, 19 recordings, 676 cells; P = 4.78e-33, Wilcoxon rank-sum test, Fig. S6D). It is plausible that the effect of vestibular lesions are indirectly caused by the suppression of vestibulo-ocular reflex (VOR) driven eye movements. We thus further determined the extent to which RSP neuronal activity is modulated with eye position during VOR resets, and found a very small fraction of RSP neurons with eye-movement correlated activity (cells with significant pupil position correlations in the dark: temporal = 8/101, nasal = 6/101, n = 3 mice, Fig. S7). These findings not only highlight the contribution of vestibular cues to AHV signalling, but also show that eye movement-related signals are not significant drivers of AHV tuning.

Since the RSP is extensively connected with visual areas (51–53), it is conceivable that, when available, optic flow information could also contribute to motion representation by AHV neurons. To test this, we recorded from AHV cells during both vestibular stimulation (passive rotation in the dark, Fig. 2A) and full-field visual motion stimulation when the animal was stationary (Fig. 2B). Since the visual motion stimulus, which was projected onto a cylinder surrounding the animal, had the same velocity profile as the rotation stimulus, it simulated the optic flow that would be experienced if mice were rotated past a static visual cue (Fig. 2B). Although visually-evoked responses were weaker than those evoked by vestibular stimulation (rotation modulation index: median [vestibular] = 0.29, IQR = [0.13 – 0.6], median [visual] = 0.19, IQR = [0.08 – 0.42], n = 120 AHV cells, P = 2.4e-7, Wilcoxon signed rank test), many AHV neurons (52/120) increased or decreased their spiking activity in response to both stimuli, and were modulated by the speed (62/120) and – to a lesser degree – the direction (22/120) of both (Fig. 2A-B).

**Fig. 2.**
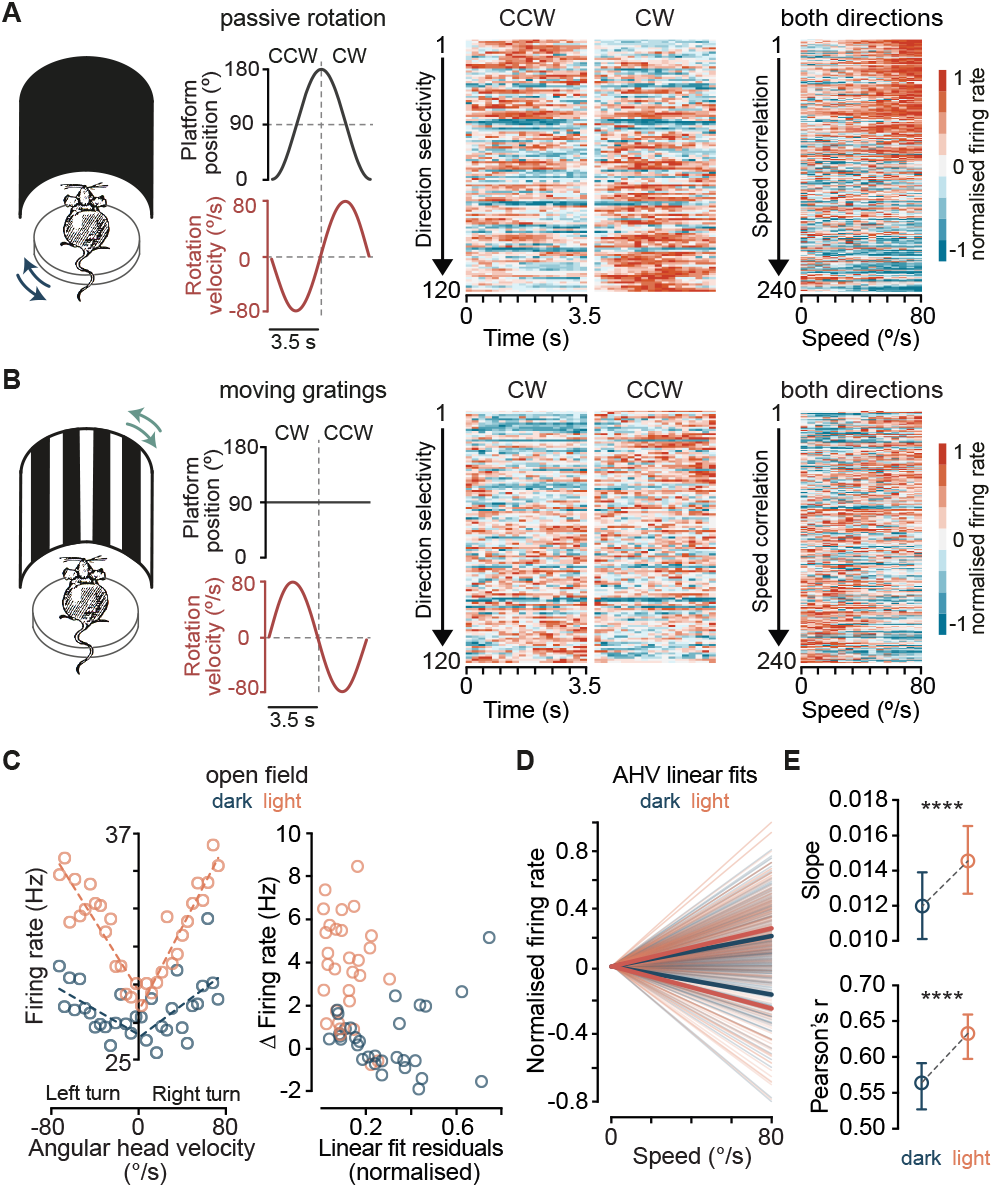
Vestibular and optic flow information converge onto AHV neurons. **(A)**Left, experimental design for vestibular stimulation. Traces show the position (black) and velocity (red) of the rotating platform. Centre, firing rate heatmaps of all recorded AHV cells during clockwise (CW) and counter-clockwise (CCW) rotations in darkness. Rows show baseline-subtracted, normalised average firing rate of all neurons sorted by their direction selectivity index (area under the direction ROC curve, see Materials and Methods). Right, baseline-subtracted, normalised average firing rate of all neurons as a function of rotation speed. Each pair of rows represents an individual neuron’s response to both directions. Neurons are sorted by the magnitude of speed correlations (Pearson’s r), averaged over the two directions. **(B)** Left, Left, the rotation platform was stationary (black trace) while a surround vertical grating was rotated with the same velocity profile as the vestibular stimulus in A (red trace). Centre and right, same as in (B) but for simulated optic flow. **(C)** Left, tuning curves of an example AHV cell recorded in the open-field in darkness and in light. Dashed lines represent linear fits. Right, for the same cell, firing rate difference between the first and all other velocity bins is plotted against the residual of the linear fit. **(D)** Population data showing linear fits for all AHV cells and for each direction in darkness and in light. Thick lines are averaged fits for either positive or negative correlations. **(E)** Summary data (median and 95 % CI, n = 272 AHV cells) showing the overall change between dark and light conditions. **** P [slope] = 2.9e-11, **** P [Pearson’s r] = 1.2e-6, Wilcoxon signed rank test.

While it is well established that visual inputs convey landmark information to head direction cells to maintain their tuning (54, 55), the contribution of vision to AHV signalling is unknown. The combined vestibular-visual activation could theoretically provide a moment-to-moment AHV signal based on both internal and external cues. In agreement with this, we observed that a significantly larger fraction of RSP cells encode AHV in presence of visual signals than in the dark in freely moving mice (dark = 192/359, light = 223/359, P = 0.02, Fisher’s exact test). Notably, addition of visual input increased the slope of the angular head velocity/spiking rate relationship and the magnitude of correlations (Fig. 2C-E), suggesting that visual signals increase the gain and improve the signal-to-noise ratio of AHV coding. Considering this increase in the dynamic range of AHV tuning when both sensory cues are available, we hypothesised that vestibular-visual integration could markedly improve the fidelity of angular velocity representation by RSP network and and generate a reliable estimate of angular self-motion.

To address this hypothesis at the behavioural and network level, we took advantage of our head-restrained paradigm which allows the separation and quantitative control of visual motion and vestibular information. We could therefore examine whether a switch to using visual flow instead of vestibular signals occurs when vision is available, or whether the two self-motion inputs are integrated for perception and neuronal encoding. First, we trained mice in a go/no-go task to discriminate between their own angular speeds, either in the dark or in presence of optic flow information (Fig. 3A-B). In complete darkness, mice could differentiate between pairs of rotation stimuli (44) that differed by at least 20 °/s in their peak speed (mean ± SEM discrimination accuracy of 5 blocks: 10°/s v. 10 °/s = 49.6 ± 3.7 % compared to 10 °/s v. 30 °/s = 74.6 ± 4.6 %, n = 5 mice, P = 0.02, one-way ANOVA with Holm-Sidak’s test, Fig. 3C). Presenting a static vertical grating (Fig. 3B), to provide optic flow in addition to vestibular stimulation, lowered the discrimination threshold from 20 °/s to 10 °/s (10 °/s v. 10 °/s compared to 10 °/s v. 20 °/s: P [vestibular] = 0.09, P [vestibular + visual] = 0.01, one-way ANOVA with Holm-Sidak’s test, Fig. 3C), and significantly increased the perceptual accuracy of self-motion (Fig. 3C, E). Notably, mice performed substantially worse in a pure visual task, which consisted of pairs of visual motion stimuli moving horizontally past the animal with the same speed profiles as the rotation stimulus pairs (Fig. 3B right, D-E). The improved self-motion discrimination performance in presence of the static visual cue is therefore not due to sole reliance on visual flow information, but rather requires the combination of vestibular and visual signals. These results demonstrate the behavioural significance of vestibular-visual combination in rodents for the first time, and is consistent with human data showing improved perception of angular self-motion with multisensory signals (56, 57).

**Fig. 3.**
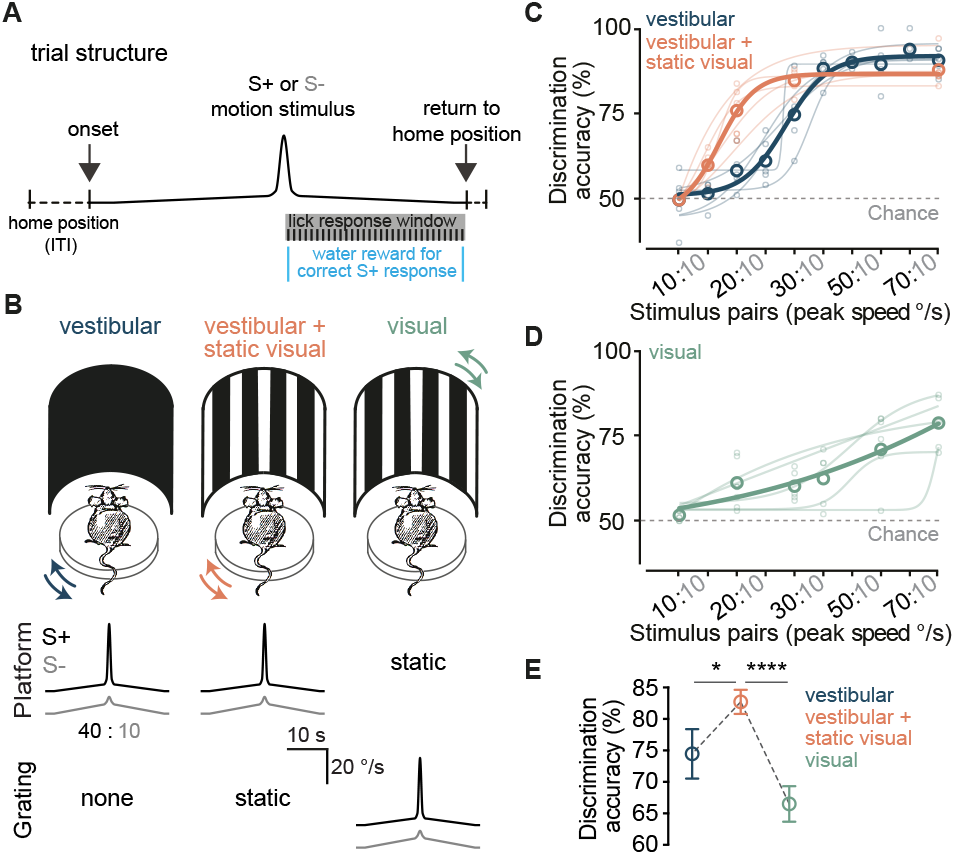
Combination of vestibular and visual cues improves perceptual accuracy of angular self-motion. **(A)** Schematic of trial structure for the go/no-go speed discrimination task. The trace illustrates a generic rotation velocity profile (motion stimulus) for an individual trial. Stimulus duration = 32.2 s, inter-trial interval (ITI) = 5 s. **(B)** Schematic of the three experimental conditions. Mice either discriminated speed of self-motion in the dark (“vestibular”, left) or in presence of a static visual stimulus (“vestibular + static visual”, centre). Under the third condition (“visual”, right), the rotation platform was kept stationary while the grating rotated to provide visual motion stimulation with the same speed profile used in the previous two conditions (centre and bottom rows). **(C)** Accuracies of self-motion speed discrimination for all stimulus pairs (n = 5 mice, average of 5 blocks). Small and large circles represent individual animals and group averages, respectively. Lines are sigmoid fits. **(D)** As in (C), but for visual motion discrimination. **(E)** Mean (± SEM) discrimination accuracies of all stimulus pairs that were tested under the three conditions (20:10, 30:10, and 80:10) in all 5 mice. * P = 0.014, **** P = 1.1e-5, one-way ANOVA with Holm-Sidak’s test.

We next sought to answer whether the activity of ensembles of RSP neurons under such multimodal condition similarly improves the fidelity of angular velocity coding, thus mirroring the psychometric data. We first employed single-cell decoding methods on AHV neurons that were tracked in the head-restrained experiments (n = 120). Decoding with ROC revealed a range of response types across the population, with some neurons discriminating the direction (clock-wise v. counter-clockwise) and/or the speed of rotations exclusively in the dark, some only in presence of a static visual stimulus, and others under both conditions (Fig. 4A-B, Fig. S8A). Despite this heterogeneity, the total number of cells that could discern the angular speed (vestibular = 36 /120, vestibular + visual = 51/120) and the average discriminability (area under ROC = AUC, Fig. S8B) increased in presence of optic flow compared to in darkness. For rotation direction, the effect of multisensory combination was only evident at the onset of motion (Fig. S8C left, first 500 ms of rotation), with no significant difference in proportion of direction-discriminating cells (entire rotation window: vestibular = 33/120, vestibular + visual = 34/120, P = 1; first 500 ms: vestibular = 5/120, vestibular + visual = 22/120, P = 8e-4, Fisher’s exact test) or discriminability between the two conditions when longer epochs of the rotation stimulus (3.5 s) were considered (Fig. S8C). We reasoned that the observed functional diversity at the level of individual AHV cells might enhance the efficiency of the population code, thus allowing the RSP ensembles to use all relevant available sensory cues to generate the most accurate representation of self-motion.

**Fig. 4.**
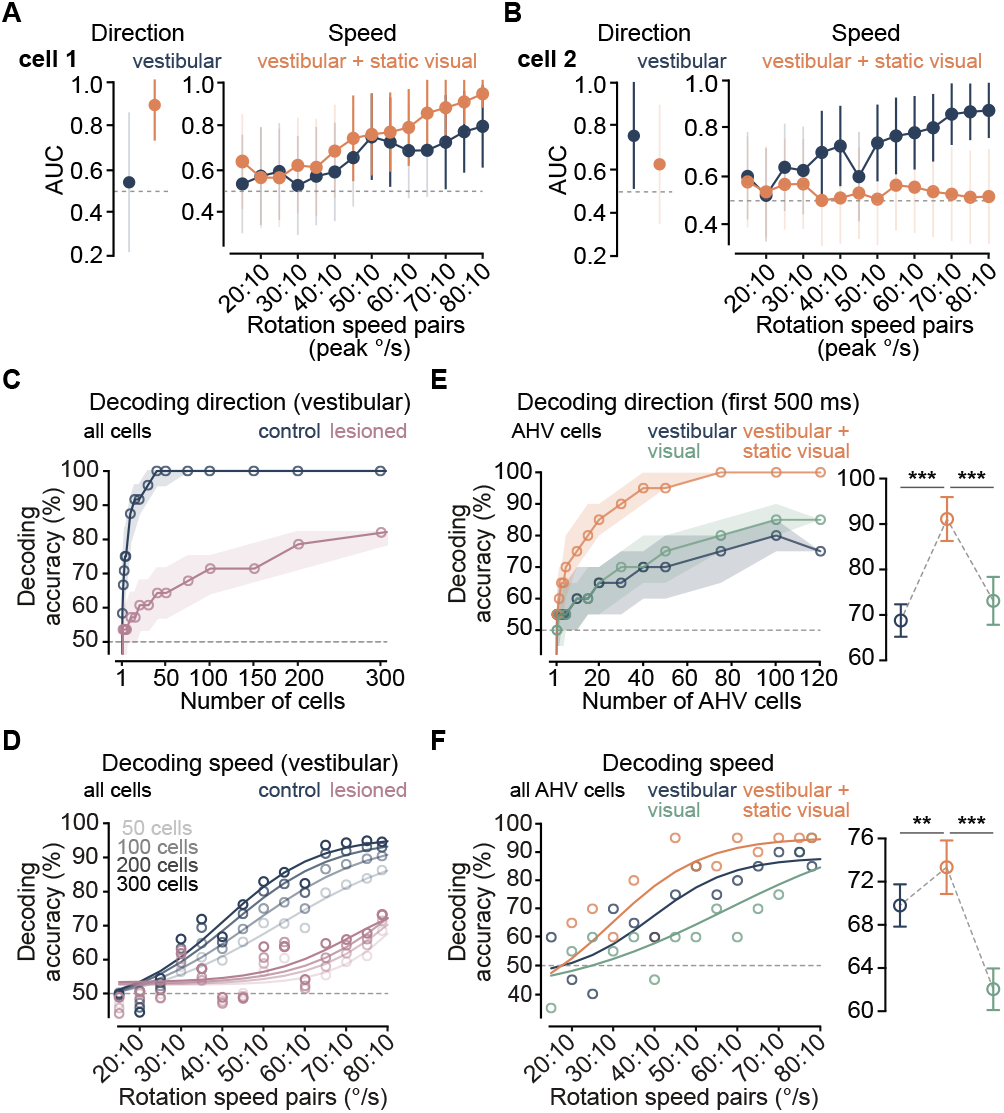
Combination of vestibular and visual cues improves decoding of angular self-motion by AHV networks in the RSP. **(A)** Left, normalised AUCs for a cell that discriminates direction of rotation in presence of visual signals (vestibular + visual), but not in the dark (vestibular). Right, normalised AUCs for speed ROC showing improved speed discrimination under the vestibular-visual condition. Transparent and solid error bars indicate 99 % CIs within and above chance level, respectively. **(B)** Similar to A, but for a different cell that discriminates direction and speed of rotation only in the dark. **(C)** LDA direction decoding accuracy (mean + IQR) in darkness as a function of population size for controls and vestibular-lesioned animals. **(D)** LDA speed decoding accuracies (10 °/s v. 15 °/s – 80°/s) in darkness with increasing population size for controls and vestibular-lesioned animals. **(E)** Left, LDA decoding accuracy (mean + IQR) for direction of self-rotation (blue and orange) and visual motion (green) as a function of AHV population size. Only the initial 500 ms of stimuli was considered. Right, mean (± SEM) decoding accuracy (5 pseudo-populations, 10 – 120 pooled neurons). *** P [vestibular v. vestibular + visual] = 4.2e-4, *** P [visual v. vestibular + visual] = 4.2e-4, one-way ANOVA with Holm-Sidak’s test. **(F)** LDA decoding accuracies for speed of self-rotation (blue and orange) and visual motion (green) using all 120 AHV cells pooled into a pseudo-population. Lines are sigmoid fits. Right, mean (± SEM) decoding accuracy from all speed pairs (5 pseudo-populations, 10 – 120 pooled neurons). ** P [vestibular v. vestibular + visual] = 0.007, *** P [visual v. vestibular + visual] = 0.0004, one-way ANOVA with Holm-Sidak’s test.

To investigate this, we used linear discriminant analysis (LDA) to decode the direction and speed of rotation from the population activity. First, by pooling all RSP neurons recorded in the dark into pseudo-populations of increasing size, we found that the decoder could reliably predict these angular velocity parameters with ensembles of fewer than 100 neurons (Fig. 4C-D). Importantly, vestibular lesions significantly reduced the accuracy of decoding both direction (mean ± SEM direction decoding accuracy: Lesioned = 66.6 ± 2.4 %, n = 4 mice; Control = 96.1 ± 1.6 %, n = 10 mice; 10 – 300 pooled neurons, P = 1.7e-9, Student’s t-test, Fig. 4C) and angular speed (mean ± SEM speed decoding accuracy: Lesioned = 55.3 ± 0.7 %, Control = 70.5 ± 2.6 %, all speed pairs, P = 0.0001, Student’s t-test, Fig. 4D). These data suggest that a small ensemble of RSP neurons can reliably report angular head velocity in the dark using vestibular input alone. Next, using the same method on all tracked AHV neurons, we compared decoding accuracies under the three experimental conditions. As predicted, adding a static vertical grating during rotation enhanced the performance of the decoder and, similar to the perceptual task, this improved performance could not be attributed to vision alone (Fig. 4E-F); however, in line with results from the ROC analysis, the improvement in decoding direction was only evident at the beginning of motion (mean ± SEM direction decoding accuracy for the entire rotation window: vestibular = 95.8 ± 2.9 %, vestibular + visual = 96.6 % ± 2.5 %, 5 pseudo-populations, 10 – 120 pooled neurons, P = 0.16, one-way ANOVA with Holm-Sidak’s test, Fig. 4E). This may be explained by the relatively slow processing time of vestibular compared to visual information (58), and the significance of multisensory integration in resolving ambiguities at the beginning of motion. Finally, we obtained similar results when using all populations of simultaneously recorded RSP neurons (Fig. S9).

## Discussion

We have provided the first quantitative analysis of sensory mechanisms underlying cortical AHV coding. Our results reveal a previously under-appreciated extensive network of AHV neurons in the RSP that is dominated by vestibular input and can reliably encode the direction and speed of head turns during both active exploration and passive motion. The AHV (and locomotion speed) cells in the RSP may be used in path integration by updating the spatial map as the animal moves through the environment, and possibly contribute to perception of self-motion. These neurons may also engage in generation and updating of the recently identified egocentric goal-vector cells in the RSP and downstream regions (38).

By demonstrating the maintenance of AHV signalling between freely moving and head-restrained conditions, we show that proprioceptive signals and efference copy of motor commands that arise from voluntary head movements and locomotion are not major drivers of retrosplenial AHV neurons. Nevertheless, considering that tuning properties of some AHV cells differed between the two conditions, motor-related signals may provide additional input to these neurons and modulate angular velocity coding during navigation. This interpretation is consistent with the observation that many vestibular neurons in the brainstem integrate proprioceptive and motor related information (59). Alternatively, differences between active and passive AHV tuning may arise from cortical posture related signals in freely moving animals (50), and/or the substantially more complex head movements during free exploration which encompass multiple planes of motion in three-dimensional space. The absence of substantial eye movement-related activity in the RSP further suggests that eye-movement-sensitive neurons in the vestibular nuclei (60), which are modulated by pupil position and the fast phase of VOR, are not the primary source of AHV tuning in the RSP. However, it is likely that brainstem neurons with integrated eye and head movement information further contribute to the ascending AHV signal (23, 24).

The prevalence of AHV cells in the RSP and the large fraction of conjunctive neurons identified here is at odds with the current network model of the mammalian head direction system, which assumes that cortical areas such as the RSP inherit a pure allocentric heading representation (61). In this hierarchical model, AHV and conjunctive cells predominantly reside in brainstem nuclei where the head direction signal is generated by integrating the AHV over time, and passed along to higher thalamocortical areas. In contrast, our data suggest that the neural substrates required for the generation and updating of the head direction signal also exist in the cortex. Interestingly, the proportion of AHV neurons in the RSP and their tuning properties resembles those described in the brainstem (11, 12). It is therefore likely that multiple cortical and subcortical regions are concurrently involved in head direction computations.

We have also shown that additional visual input to the RSP increases the gain and signal-to-noise ratio of individual AHV neurons and the fidelity of encoding the direction and speed of head turns at the network level. This is in line with our behavioural data, and previous work in primates (17–21), showing improved perception of self-motion when visual cues are present. Therefore, while vestibular input is critical for encoding AHV, visual input has a gain-of-function role in self-motion signalling during navigation. The contribution of vision in this process most likely consists of the combined effect of optic-flow, which provides additional motion information, luminance (62), which can modulate AHV tuning, and the improved gain of gaze-stabilising eye movements when VOR and optokinetic reflexes work together (63). We thus propose that the significance of visual input to the RSP – and likely to other parts of the head direction network – extends beyond allocentric representation of spatial landmarks (32, 45-48). Instead, both allothetic and idiothetic visual information can be used to maintain accurate spatial orientation.

These data provide a mechanistic explanation of turn-specific modulations of neural activity during route running observed in the RSP (31) and beyond (64–66). We also highlight a critical role of the vestibular sense in cortical processing, which despite its vital role in spatial cognition has received relatively little attention. The robust representation of vestibular stimuli observed here complements and extends recent work on vestibular-evoked responses in the rodent cortical network (44, 62, 67). AHV neurons in the RSP may belong to the same population of cells that were previously shown to provide head motion information to the primary visual cortex (44), and may also be the source of head motion signals in other cortical areas that are directly connected to the RSP, such as postsubiculum (68) and the entorhinal cortex (69). The pathways that provide AHV information to the RSP are yet to be described but one possible route could involve the ascending thalamocortical head direction network. However, there is no evidence of substantial AHV coding in the anterior thalamic nuclei. Alternatively, these signals may arise from the posterior vestibular pathways and reach the RSP indirectly via other associative cortical regions (24). Irrespective of the source of these inputs, there appears to be a significant egocentric component to motion signalling in the RSP, indicating that the this area can combine both environmental features (34, 35, 38) and correlates of movement into a coherent egocentric spatial map.

## Materials and Methods

### Animals

A total of 18 adult (8 – 14 weeks old, 25 – 35 g) male C57BL/6 mice were used in accordance with the UK Home Office regulations (Act of 1986, Scientific Procedures) and the Animal Welfare and Ethical Review Body (AWERB). Mice were maintained on a 12-hr light/dark cycle and single housed after surgical procedures.

### Surgical Procedures

All surgical procedures were carried out under anaesthesia either with isoflurane (2 % – 5 %) or with a mixture of fentanyl (0.05 mg/kg), midazolam (5.0 mg/kg), and medetomidine (0.5 mg/kg) in saline solution (0.9 %, i.p.). Mice were then fixed in a stereotaxic frame and eyes were protected with a lubricating eye gel (Lubrithal, Dechra). Body temperature (37 – 38 °C) and anaesthesia level was maintained throughout the procedure. Carprofen (5 mg/kg, s.c.) was administered for postoperative analgesia and, where appropriate, injectable anaesthetics were reversed with a mixture of naloxone (1.2 mg/kg), flumazenil (0.5 mg/kg) and atipamezole (2.5 mg/kg) in saline solution (0.9 %, i.p.).

For probe recordings, a custom-built titanium head fixation implant was affixed to the skull using a cyanoacrylate-based adhesive (Histoacryl, Braun) and dental cement (Simplex Rapid, Kemdent). A 1 mm craniotomy was performed with a 0.3 mm burr dental drill (Osada Electric, Japan) over the right RSP (AP: -3.40 to -2.20 mm, ML: 0.55 to 1.10 mm). For acute recordings, the craniotomy was performed 3 – 24 hours in advance and sealed with a removable silicone sealant (Kwik-Cast, World Precision Instrument). Chronic probe implants were fixed to the skull using a light-cured dental cement (3M). In addition, a gold pin was inserted inside a craniotomy rostral to the Bregma, secured to the skull, and attached to the ground wire. Neuropixels implants were additionally secured inside a custom-built 3D printed enclosure (38).

For bilateral vestibular lesions, an incision was made behind each ear and muscles covering the temporal bone were bluntly dissected to expose the posterior and horizontal semi-circular canals. Canal bones were then thinned with a 0.3 mm burr dental drill until punctured and a microfiber needle was inserted into each canal to deliver 50 µl of kanamycin (50 mg/ml) in distilled water. After approximately 5 minutes, the solution was removed with a sterile cotton tip and the wound was closed using a cyanoacrylate-based adhesive.

### Recording setups

#### Open-field experiments

Open-field recordings were performed on an elevated 92-cm diameter circular arena located inside a 140 x 140 x 160 cm sound-proof enclosure (38). A surrounding hectagonal black wall (45 cm high from the arena surface) was placed at approximately 10-cm distance from its circumference. Six infrared LEDs (TV6700, Abus) were positioned on enclosure walls to provide illumination. Mice were allowed to freely explore while their behaviour was monitored at 40 frames per second (fps) with a near-IR camera (acA1300-60gmNIR, Basler) centred above the arena. During light conditions, the background luminance was set at 11.5 lux using a projector (BenQ) pointing at a translucent overhead screen (Xerox). Video recordings were controlled by a custom software written in Lab-VIEW (2015 64-bit, National Instruments) and Mantis software (mantis64.com). A PCIe-6351 board (National Instruments) was used to control triggering of each camera frame and for synchronisation. A Neuronexus commutator or a custom-made rotary joint (adapted from Doric AHRJ-OE_PT_AH_12_HDMI) was used to prevent twisting of probe cables during recording. To correct for any potential drift between camera triggers and probe recording, a micro-controller (Arduino Leonardo) was used to continuously deliver logic pulses of varying time durations (randomly sampled from a uniform distribution).

All open-field sessions consisted of three experimental conditions, each lasting 20 – 30 minutes. Recordings were first made in light (light1) where two side-by-side white cue cards (88 x 60 cm) were attached inside of the otherwise featureless black wall. Further recordings were made in darkness, and again in a second identical light condition (light2). At the completion of each open-field recording session, mice were taken to the head-restrained passive rotation apparatus. The two recordings always occurred on the same day with a maximum of 5 hours in between.

#### Passive rotation experiments

Mice were head-fixed and restrained in a custom-made tube mounted onto a recording platform that was attached to the rotation motor (RV120CC, Newport Corporation). The animal’s head was positioned such that the axis joining the ears was parallel to the horizontal plane of motion, with vestibular apparatuses positioned around the axis of rotation (44). Prior to recordings, mice were habituated to head fixation and rotation for 2 – 3 days with each habituation session lasting 15 – 30 minutes. Throughout habituation mice were given a sweetened condensed milk reward.

Visual stimuli were presented with a custom-built 3D surround projection system made of two laser projectors (MP-CL1A, Sony) and a cylindrical rear-projection screen (radius = 10 cm, height = 15 cm, central angle = 300°). The cylindrical screen was positioned concentrically with the rotation motor such that the animal’s head was positioned at its centre. Control of visual stimuli, alignment of images from the two projectors, image warping, edge blending and luminance correction were all implemented using custom algorithms written in Bonsai (70). The animal’s field of view was restricted with a pair of blinkers (azimuth = 120°, altitude = 50°) to prevent viewing beyond the screen boundaries during rotation. The visual stimulus consisted of either stationary or rotating full-field vertical square-wave gratings, with a mean luminance of 16 lux and a spatial frequency of 0.04 cpd. Horizontal rotation of the recording platform, or the vertical grating, was achieved using custom-written routines in Igor Pro in combination with NeuroMatic (71). Rotation stimulus consisted of a full sinusoidal period (7 s), flanked by sinusoidal ramps of the same period, and reached a maximum displacement of 90° and a maximum velocity of 80 °/s in each direction (CW or CCW, Fig. 1C). Each rotation stimulus was separated by a stationary period (10 – 30 s) and repeated 10 – 15 times under each experimental condition. Waveforms used to rotate the platform and the surround visual stimulus were identical. To block any spatial auditory cues from the rotation motor or the environment, a masking white noise was played throughout the recording using a piezoelectric miniature speaker (Sonitron) mounted behind the animal on the platform. Synchronisation with the probe recording was achieved by logic signals via an ITC-18 board. In addition, a photodiode (PDA100A-EC, Thorlab), placed out of the field of view behind the projection screen, was used to precisely record the onset of visual stimuli.

Each recording session consisted of three experimental conditions (Fig. 3B): 1) rotation of the mouse in complete darkness (“vestibular”), 2) rotation of the mouse in presence of the static surround vertical grating (“vestibular + visual”), and 3) rotation of the vertical grating while the platform was motionless, producing simulated optic flow (“visual”). Trials from these three conditions were distributed pseudo-randomly throughout the recording session.

### Single-unit recordings

Three mice were chronically implanted in the right RSP with the Neuropixels (phase3A, option 1 and 2, 384 channels) (72) and 2 with the Neuronexus probe (Poly2, 32 channels). All acute recordings were made with the Neuropixels probe (phase3A, option 1, 2, and 3). Prior to each insertion, the probe shank was coated with DiI (1 mM, Thermo Fisher Scientific) to allow post-hoc histological reconstruction of the probe track (Fig. S1A).Dura was left intact and the craniotomy was covered with saline while the probe was lowered at a speed of 2 µm/s to the desired depth using the stereotaxic manipulator (Kopf Instruments) or micromanipulators (SM-8, Luigs and Neumann). For acute recordings, the craniotomy and the Ag/AgCl ground pellet (1 × 2.5 mm, World Precision Instruments) were then submerged in a 2 % agarose (in saline) solution and the probe was allowed to settle for at least 15 minutes before recording started. For chronic implants, the craniotomy was sealed with a silicone sealant (Kwik-Cast, World Precision Instrument). Mice with chronic implants were allowed a minimum of 5 days of recovery from surgery before recordings started and underwent 2 – 3 recording sessions. Successive recording sessions were separated by a minimum time interval of 3 days. At the end of recordings, probes were retracted from the brain, submerged in 1 % Tergazyme (in distilled water, Alconox) for at least an hour, and further rinsed with distilled water. For chronic implants, the brain underwent fixation by transcardial perfusion before the probe was retracted.

Extracellular potentials were acquired using an FPGA card (KC705, Xilinx) or the Whisper system (Janelia Applied Physics and Instrumentation Group), and spikeGLX was used as the data acquisition software (https://github.com/billkarsh/SpikeGLX, Janelia Research Campus). For Neuronexus recordings, data were amplified with a gain of 200, filtered at 0.1 – 10000 Hz, and digitised to 16 bits at 25 kHz. For Neuropixels recordings, signals were amplified with a gain of 500, high-pass filtered at 300 Hz, and digitised to 16 bits at 30 kHz.

To isolate single units, data were band-pass filtered (300 – 5000 Hz), and the median of each channel was subtracted to remove any baseline offset. Correlated sources of noise were then removed by common (median) average referencing in blocks of 77 (Neuropixels) or 32 (Neuronexus) channels. Automated spike sorting was carried out using Kilosort (http://github.com/cortex-lab/KiloSort, (73)) or KiloSort2 software (http://github.com/MouseLand/Kilosort) with further manual curation of the isolated units using Phy (https://github.com/cortex-lab/phy). Spike sorting parameters are detailed in Tables S1 – S2. Units were excluded if they had < 900 spikes (minimum firing rate = 0.2 Hz), absolute refractory period of < 1 ms, significant amplitude drift across the recording session, and atypical waveforms (noise-like or only positive potentials).

### Anatomical assignment of single units

At the end of the recording, animals were deeply anaesthetised with 120 mg/kg pentobarbitone (Euthatal, Merial Animal Health) and transcardially perfused with chilled heparinised phosphate buffer (10 U/ml in PB 0.1 M) and 4 % paraformaldehyde (PFA) in PB (0.1 M). Following another 24 hours of fixation in 4% PFA, the brain was extracted and either embedded in 4 % agar for automated whole-brain imaging or cut coronally into 100 µm sections using a vibratome (Microm HM 650V, Thermo Scientific). Coronal sections were mounted in a medium containing DAPI (Santa Cruz Biotechnology) and imaged with an epifluorescence microscope (Axio Imager 2, Zeiss). Automated whole brain imaging was performed using serial two-photon tomography. Acquired images were either warped manually into matched sections in the Allen Mouse Brain Atlas (74), or by using a validated registration pipeline implemented within the cellfinder software (https://github.com/brainglobe/cellfinder, (75, 76)). The probe track was identified by DiI fluorescence and reconstructed manually using drawing tools in Fiji (77). The end of the track was determined as the point where DiI fluorescence was no longer visible. This tip location closely matched the probe depth acquired from manipulator readings. By measuring distances from the probe tip and using the geometrical configuration of the probe, the anatomical location of all recording sites along the track was determined. To increase accuracy, these measurements were further compared with electrophysiological signatures along the probe track (i.e. the lack of activity in the white matter or the change in signal-to-noise ratio at the interface of brain and solution), and adjusted accordingly. To identify the location of isolated units along the probe, an average waveform (first 100 spikes) was generated and the recording channel with the largest negative peak was determined.

### Single-unit tracking

To identify the same units between freely moving and head-fixed recordings, we developed a single-unit matching pipeline to identify same units between the two conditions. Spike sorting was carried out separately on the two recordings, and for each isolated unit in the head-fixed data (here defined as “Fix”), candidate matches in the freely moving recording (here defined as “Free”) was chosen from either eight neighbouring channels when using the Neuropixels (four in each direction, maximum 40-µm distance), or four when using the Neuronexus probe (two in each direction, maximum 50-µm distance). A similarity score was then calculated for all “Free” units on these neighbouring channels based on three categories of similarity metrics derived from the average spike waveform, spiking rate, and inter-spike interval (ISI) distributions:

#### Spike rate/ISI

For each “Fix” unit and its candidate “Free” matching units, the average spiking rate and the ISI histogram was obtained from the entire duration of the experiment. The following three comparative parameters were then calculated: 1) Spiking frequency ratio, defined as the ratio of the average spiking rate between the “Fix” and candidate “Free” units. Ratios above 1 were rectified. 2) ISI histogram correlation, defined as the Pearson’s r of “Fix” versus “Free” ISI histograms. 3) ISI histogram intersection, defined as the intersection ratio between “Fix” and “Free” ISI histograms calculated as:

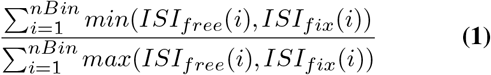

Where ISI_fix_ and ISI_free_ are the ISI histograms of the “Fix” and the candidate “Free” matching unit, respectively.

#### Local spike features

Four parameters were used to quantify the difference between specific features of the average “Fix” and “Free” waveforms, including the difference in: 1) the height of the 2nd peak, defined as F(B_fix_, B_free_), 2) trough-to-peak duration, defined as F(C_fix_, C_free_), 3) trough-to-peak amplitude, defined as F(B_fix_ – D_fix_, B_free_ – D_free_), 4) peak-to-trough amplitude, defined as F(A_fix_ – D_fix_, A_free_ – D_free_).

Where A is the maximum amplitude of the first positive peak preceding the trough; B the maximum amplitude of the second positive peak following the trough; C the time between the minimum of the initial trough and the maximum of the following peak; D the minimum amplitude of the initial trough; and F(x,y) the proportional difference function given by the following equation:

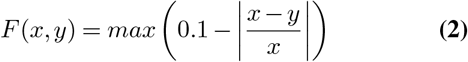

#### Global spike features

The following three parameters were calculated to compare the entire average spike waveform (4 ms window, from -160 µs to 240 µs relative to the trough) between the “Fix” and the candidate “Free” unit: 1) spike waveform histogram intersection, defined as the intersection ratio between “Fix” and “Free” histograms of the amplitude of the average spike waveform (3-µV bins) calculated as in Eq. (1). To quantify the temporal similarities between the two waveforms, the following three parameters were calculated by first applying dynamic time warping (DTW, (78)) to the average spike waveforms using the fastdtw package (http://github.com/slaypni/fastdtw). 2) spike waveform difference, defined as the maximum Euclidean distance between the dynamic time warped “Fix” and “Free” average spike waveforms (normalised by the peak-to-through amplitude). 3) spike waveform correlation, defined as the Pearson’s r of “Fix” versus “Free” dynamic time warped waveforms. 4) DTW scale factor, defined as the ratio of the length of the original “Fix” average spike waveform to the length of the signal after DTW, so as to penalise excessive signal warping.

For each category of similarity metrics described above, the Euclidean distance between all parameters was calculated, thus generating three similarity metrics. Empirical weights were then applied to each metric (W_local_ = 0.25, W_global_ = 1, W_rate/ISI_ = 1) and a final similarity score was calculated by summing the three weighted metrics. For each “Fix” unit, the “Free” unit with the largest similarity score was identified as the best potential match. If a given “Free” unit was identified as the best potential match of more than one “Fix” unit, it was only assigned to the one with which it had the highest similarity score. At the final step, the best potential match was accepted as the matching “Free” unit if the following thresholds were met: ISI histogram correlation > 0.65, all four local spike features > 0.5, spike waveform correlation > 0.95, spike waveform difference < 0.3. Empirical weights and thresholds were decided by visual inspection of a subset of data using a graphical user interface.

#### Single-unit classification

K-means clustering (with k = 2) was used to classify the isolated units as wide or narrow spiking based on features of their average waveform at the recording site with the largest amplitude. Two metrics were used for this classification: the height of the second positive peak relative to the initial trough, and the time between the minimum of the initial trough and the maximum of the following peak.

To investigate whether wide- and narrow-spiking cell classes correspond to excitatory and inhibitory cell types, respectively, we next performed cross-correlogram (CCG) analysis to identify putative monosynaptic connections between neurons (79). Excitatory and inhibitory connections were identified as short-latency and short-duration peaks or troughs in the CCGs. The baseline firing rate of each 0.5-ms CCG bin (−50 to 50 ms) was first determined by applying a Fourier smoothing with a frequency cut-off at 5 kHz. A confidence band was then calculated from the 0.0001 – 99.9999 percentile of the cumulative Poisson distribution at each time bin, and used as the statistical threshold for detecting outliers at 1.5 to 4 ms time window from the centre bin. A significant peak, indicating an excitatory interaction, exceeded the upper limit of the confidence band in at least two bins. A significant trough, indicating an inhibitory interaction, was below the lower limit of the confidence band in at least three bins. Cell type assignment using K-means clustering was then evaluated for the subset of units identified as excitatory and inhibitory based on CCG interactions, and the two were found to be largely overlapping (93.3 % overlap).

### Pupil tracking

The right pupil was recorded at 40 fps using a near-IR camera (acA640-750um, Basler AG) and a custom program written in C++ based on the OpenCV and Pylon 5 libraries (https://www.baslerweb.com/en/sales-support/downloads/software-downloads, (44)). An ITC-18 interface board (InstruTECH, Heka Elektronik) and Igor Pro were used to control camera triggering and synchronisation with rotation stimuli. Illumination was provided by an array of infrared LEDs (Kingbright, 940 nm) and the pupil was focused using two lenses (f1 = 25 mm, f2 = 75 mm, Thorlabs). Pupil movements were then tracked offline using DeepLabCut (DLC, (80)). A pre-trained ResNet-50 network was trained on 240 frames from 6 video recordings (40 frames from each) with 500000 training iterations, reaching a final loss of 0.0007 and a training and testing error of 1.7 and 4.1 pixels, respectively. To identify rapid eye movement events, the differential (central derivative) of the horizontal eye position over time was calculated and a median filter (50-ms window) applied to attenuate noise. Outliers were removed by using a threshold cut-off of 20 mm/s and a Lowess smoothing (span = 0.01) was applied to subtract low frequency fluctuations. Rapid eye movement events were then detected using a threshold of 2.5 × SD. For each isolated unit, the number of spikes that occurred during nasal or temporal eye movement events was obtained (from 75 ms before event onset to 250 ms after) and the average spiking rate histograms (25-ms bins) constructed. Neurons were considered to be modulated by eye movements if their average spiking rate histogram was significantly correlated (Pearson’s r test) with the average amplitude of the eye movement events either at nasal or temporal direction.

### Vestibular lesion assessment

Mice were allowed 4 – 11 days to fully recover from the vestibular lesion surgery before behavioural and single-unit recordings were made. All lesioned mice showed body curling when held by the base of the tail. Further vestibular deficits were quantified by recording the animals’ trajectory while exploring a 39-cm diameter circular arena in the dark using Raspberry Pi 1B, a Pi NoIR camera (30 fps) and a custom-written routine in Python (http://github.com/SainsburyWellcomeCentre/Pyper, (44)). The animal’s trajectory was recorded both before the lesion surgery and after full recovery. The Cartesian coordinates of the centroid of the mouse were determined for each frame, and turning angle was quantified by calculating the change in trajectory in the horizontal plane using coordinates from three consecutive frames. The distribution of turn angles over a set distance was then compared between pre- and post-lesion recordings.

### Rotation discrimination task

All behavioural experiments were performed on 6-week old mice. Animals were allowed to recover for at least two days from head-plate implant surgery before training began. They were then given restricted access to water for another two days and habituated to head-fixation on the rotating platform. The head was positioned over the centre of the axis of rotation as above and a water reward port was positioned in front of the animal. Mounted on the same platform was a roofed Perspex cylinder (radius =12 cm) that completely surrounded the animal. The outside of the cylinder was coated with a black projection film onto which the visual stimulus was projected. The entire rotation platform was housed inside an acoustically and optically isolated chamber.

Starting with the “vestibular” training, mice were rotated 90° clockwise in the dark over a period of 32.2 s with a peak velocity of 80 °/s (S+ stimulus, Fig. 3A-B), and for at least 100 trials (inter-trial interval = 5 s). Licking for two or more 250-ms time bins after the peak velocity (16.1 s from rotation onset) yielded a water reward of approximately 5 µl, and was scored as a correct S+ response. Once mice demonstrated licking in more than 80 % of S+ trials for at least 100 trials, the S-stimulus was introduced. The S-stimulus had a peak velocity of 10 °/s, identical onset and offset times and area under the curve to the S+ stimulus (Fig. 3B), and was not rewarded. Licking for fewer than two 250-ms bins after the peak velocity of S-stimulus was recorded as a correct S-response. Mice were pseudo-randomly presented with 10 S+ and 10 S-trials in blocks of 20 trials, and performed 5 – 15 blocks per day. The percentage of correct responses was determined for each block, and discrimination accuracy was measured as the sum of the percentage of correct S+ and S-trials, divided by two. Once mice reached a criterion of 80 % discrimination accuracy on average for at least 5 consecutive blocks, and performed a minimum of 20 blocks, the S+ stimulus peak was reduced incrementally by 5 – 10 °/s every 5 blocks until the S+ and S-stimuli were identical.

Next, mice were tested in the “visual” experimental condition similar to the recording paradigm above. The platform remained motionless and a vertical grating was presented on to the projection film using the laser projector and moved around the mouse. Here, the projector was mounted on a post attached to the rotation motor, and the grating (azimuth = 56°, altitude = 50°) was physically rotated 90° in the CCW direction starting at 45° to the right of the head’s midline. Mice were trained to discriminate between pairs of rotating visual stimuli, with speed profiles identical to S+ and S-vestibular stimuli described above. After performing at least 20 blocks on the stimulus pair consisting of an 80 °/s peak velocity (S+) and a 10 °/s peak velocity (S-), the peak speed of the S+ stimulus was again reduced by 5 – 10 °/s every 5 blocks until the two stimuli were identical. Finally, under the “vestibular + visual” condition, the visual stimulus remained static and the mouse was rotated as in the “vestibular” condition.

To compare performance between experimental conditions, for each condition, discrimination accuracies from all mice and for the three stimulus pairs that were tested under all experiments (S+ peak = 20 °/s, 30 °/s, and 80 °/s) were grouped. Group means were then compared using one-way ANOVA followed by the Holm-Sidak multiple comparisons test.

### Analysis

#### Open-field analysis

Behavioural variables were extracted from video frames using DLC. A pre-trained ResNet-50 network was trained on 1190 frames from 8 video recordings (40 – 70 frames from each) with 700000 training iterations, reaching a final loss of 0.0009 and a training and testing error of 1.9 and 2.8 pixels, respectively. Head direction and AHV were calculated from tracked ear positions (horizontal plane, top view). Locomotion speed was calculated from tracked body position (Fig. 1A). Cells with average firing rate < 0.5 Hz over the entire experiment duration were excluded from the analysis. For the analysis of head direction tuning, recording epochs in which the animal’s speed was below 1.5 cm/s were excluded. This speed cut-off was not used for the analysis of speed or AHV tuning. Tuning curves were computed separately for each experimental condition (light1, dark, light2) with the same recording length used across conditions.

Head direction was defined as the angle between the horizontal axis and the line perpendicular to the axis joining the ears. To determine head direction and AHV tuning, first head direction time series was smoothed with a sliding mean filter (50-ms width). For each neuron, the average firing rate as a function of head direction, binned at 6°, was then calculated as the total number of spikes divided by the total time that the animal’s head occupied each head direction bin. The mean Rayleigh vector length of the head direction tuning curve was computed and the Rayleigh test was performed to assess non-uniformity using pycircstat (http://github.com/circstat/pycircstat). Neurons were classified as head direction tuned if under both light conditions (light1 and light2), the distribution of their firing rate as a function of head direction was statistically non-uniform (Rayleigh P value < 0.01) and their Rayleigh vector length was > 99th percentile of the mean vector length in the null distribution. The null distribution was determined by a shuffling procedure based on all recorded neurons with 1000 permutations performed for each cell. For each permutation, the entire sequence of spikes was shifted by a random amount between 20 s and the total duration of recording minus 20 s, and the mean vector length calculated from the shuffled head direction tuning curve. The 99th percentile threshold came to 0.19, consistent with previous reports in mice (81).

To construct AHV tuning curves, for each time point on the smoothed head direction times series, the angular velocity was calculated by determining the first derivative over a 200-ms time window starting from that time point. The firing rate was then plotted as a function of angular velocity binned at 6°/s. To allow comparison with passive rotation experiments, we used a maximum angular velocity of 80 °/s to construct the AHV tuning curves. Positive velocities were assigned to right and negative velocities to left head turns. For each cell, two AHV scores were calculated for right and left head turns, defined as the magnitude of correlation (absolute Pearson’s r) between the firing rate and the angular velocity. AHV slopes were calculated by applying a linear fit to the right-turn and left-turn firing rate-AHV functions. Cells were classified as AHV tuned if either of the AHV scores was > 95th percentile of the null distribution. The null distribution was based on each cell and obtained by a similar shuffling procedure as above with 1000 permutations. To compare the AHV tuning curves between light and dark conditions, neurons that met the above criterion during either dark or light1 session were classified as AHV cells.

To identify cells tuned to linear locomotion speed, the position of the mouse in each video frame was estimated in Cartesian coordinates and the instantaneous locomotion speed calculated as the distance travelled between two consecutive frames divided by the camera sampling rate (25 ms). Locomotion speed time series was then smoothed with a sliding mean filter (50 ms width) and for each cell, the average firing rate as a function of speed, binned at 1 cm/s, was calculated as the ratio between spike counts and the time spent in each speed bin. Locomotion speeds above 20 cm/s were excluded from the analysis due to low sample sizes at high speeds. Neurons were classified as speed tuned if the magnitude of Pearson’s r between the cell’s firing rate and the animal’s speed was > 95th percentile of the null distribution, generated for each cell by a shuffling procedure as described above with 1000 permutations.

#### Rotation analysis

The analysis procedures outlined below were applied similarly to data obtained under all three experimental conditions, thus the term “rotation” refers to both “vestibular” and “visual” stimuli. To construct rotation velocity tuning curves, for each neuron, the average firing rate as a function of rotation velocity, binned at 5 °/s, was calculated as the trial-averaged number of spikes divided by the total duration of each velocity bin. Two rotation velocity scores (CW and CCW) were calculated for each neuron, defined as the magnitude of correlation (absolute Pearson’s r) between the trial-averaged firing rate and the rotation velocity. Neurons were defined as modulated by rotation velocity if either of the rotation velocity scores was >95th of the null distribution. The null distribution was based on each cell and obtained by shuffling the firing rates across rotation velocity bins with 1000 permutations.

Stimulus-evoked responses were defined as significant increases or decreases in the average firing rate during either CW or CCW rotations as compared to the preceding stationary period of the same duration. Direction-modulated cells were defined as those with a significant difference between CW and CCW firing rates. In both cases, statistical significance was determined with the Wilcoxon signed-rank test. Rotation modulation index was defined as:

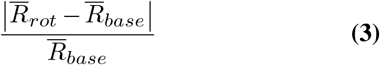

Where 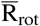 and 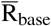 are trial-averaged (10 – 15 trials) firing rates during either CW or CCW rotations and during the stationary period, respectively.

Direction selectivity index (Fig. 2A-B) was defined as the area under the receiver operating characteristic (ROC) curve that compared the distribution of CW versus CCW firing rates. These values ranged between 0 and 1 and not rectified (see ROC analysis).

To construct the firing rate heatmaps (Fig. 2A-B, Fig. S6D), trial-averaged firing rate of each cell was calculated, binned either at 200 ms (temporal heatmaps) or 5 °/s (speed heatmaps), and normalised to the peak. The median firing rate of the preceding stationary period was then subtracted. For speed heatmaps, firing rates at CW and CCW directions were calculated separately and presented on the same plot in two consecutive rows.

To compare angular velocity tuning of neurons tracked between freely moving and head-fixed recordings, for each neuron, the average firing rate at each AHV bin (“active”) was plotted against the average firing rate at the corresponding rotation velocity bin (“passive”, Fig. 1E). Cells were defined as similarly tuned if the correlation (Pearson’s r) between the “active” and “passive” firing rates was > 95th percentile of the null distribution, computed for each cell by shuffling firing rates across AHV bins with 1000 permutations.

#### ROC analysis

We performed ROC analyses using the pROC package of R (82) to determine whether individual neurons could discriminate the direction (CW v. CCW, direction discrimination) or the speed of rotations (10 °/s v. 15 °/s – 80 °/s, speed discrimination). To construct the speed ROC curves for each neuron, the firing rate as a function of rotation speed (absolute velocity), binned at 5 °/s, was first calculated for each trial as the ratio between spike counts and the time spent in each speed bin. Speed ROC curves were then calculated by comparing the firing rate distribution of the speed bin that peaked at 10 °/s (5 °/s – 10 °/s bin) versus the successive fourteen speed bins, peaking at 15 °/s to 80 °/s. Direction ROC curves compared the firing rate distribution of CW versus CCW rotations. We computed the area under the ROC curve (AUC) as a measure of neuron’s direction and speed discriminability. AUC values lower than 0.5 were rectified, so that all AUCs were between 0.5 and 1. To test whether discriminability was significantly higher than expected by chance, for each cell, we computed the 99 % CI of the AUC values using the Delong method (83). The AUC was considered significant if the lower bound of this 99 % CI was > 0.5 (chance).

#### Population decoding

Linear discriminant analysis (LDA) was used to quantify how effectively population of RSP neurons encode direction and speed of angular motion under the three experimental conditions. This analysis was performed with the scikit-learn module (84) using Eigenvalue decomposition and applying shrinkage. LDA was used to quantify how well RSP populations could discriminate between rotation directions (CW v. CCW) or between pairs of rotation speed calculated from the firing rate-speed tuning function described above (speed bin peaking at 10 °/s v. speed bins peaking at 15 °/s – 80 °/s). For each neuron, we selected 10 trials per stimulus condition, resulting in a total of 60 trials for all three experimental conditions (“vestibular”, “vestibular + visual”, “visual”). We applied LDA to decode the direction and the speed of each trial from the firing rates of all neurons in the population, using a balanced cross-validation method for training and testing. When decoding was performed on all six conditions (2 stimulus conditions × 3 experimental conditions, Fig. 4E-F), 54 trials were used for training the decoder and 6 for testing, with a different set of trials used each time for testing. Similarly, when decoding was performed on 2 conditions (2 stimulus conditions × 1 experimental condition, Fig. 4C-D), 18 trials were used for training and the remaining 2 for testing. Before training, we z-scored all firing rates for each neuron, with no pre-selection applied based on neuron’s responses to stimuli or inhibitory/excitatory cell type assignments. Decoding accuracy was computed as the number of correctly decoded trials divided by the total number of correctly and incorrectly decoded trials. To create pseudo-populations, trial simultaneity among neurons was allocated between trials of the same stimulus and experimental condition. A bootstrap procedure was used to create pseudo-populations of increasing size and to calculate the mean decoding accuracy in each pseudo-population (Fig. 4C-F). RSP neurons from all recording sessions (or tracked AHV neurons) were sampled with replacement to generate 1000 bootstrap samples with the same number of neurons. LDA was then performed on each bootstrap sample, generating 1000 measures of decoding accuracy, and the mean decoding accuracy was computed across all bootstrap measures. To test whether the performance of the decoder improved under “vestibular + visual” compared to “vestibular” condition, for each experimental condition, mean decoding accuracies obtained from pseudo-populations of different sizes were grouped and group means compared using one-way ANOVA followed by the Holm-Sidak multiple comparisons test. To obtain a single measure of speed decoding accuracy for each pseudo-population, accuracies from all speed pairs were averaged.

The same decoding method was used on real RSP populations composed of simultaneously recorded neurons. Simultaneous populations containing a minimum of 10 or 20 neurons were used for direction and speed decoding, respectively. These thresholds were obtained from the pseudo-population analysis described above, where we determined the minimum population size required to achieve a mean decoding accuracy of above 70 % under the “vestibular” condition.

#### Statistics

For data analysis, we developed open-source pipelines in Python 3.6 using numpy (85), scipy (86), pandas (87), matplotlib (88), and seaborn (89) libraries, in addition to other packages described above. The open-field analysis code is available at: http://github.com/adamltyson/opendirection. The rest of single-unit analysis pipeline including single-unit classification, single-unit tracking, rotation and eye-movement analysis, and comparison of freely moving versus head-fixed data is available at: http://github.com/RichardFav/spikeGUI. Statistical comparisons were performed using the stats package in R or GraphPad Prism. Unless stated otherwise, statistical significance is defined as P < 0.05. Data were subjected to a test of normality with the Agostino Pearson test or Shapiro-Wilk test, and non-parametric statistics were applied to data with non-normal distributions. Statistical details are provided in the main text or in figure legends.

## Data Availability

Data will be made available upon request from the corresponding author. Data analysis codes are open-source and available via links provided within the Materials and Methods section.

## Acknowledgements

We would like to thank Jasper Poort and Pter Znamenskiy for useful discussions. The authors are grateful to Charly Rousseau for his contribution to data acquisition/analysis pipelines and Gonçalo Lopes for the development of 3D visual stimulus presentation in Bonsai. We also thank Molly Strom, Rob Campbell, the SWC Neurobiological Research Facility and FabLabs for technical support. This research was funded by the Sainsbury Wellcome Centre Core Grant from the Gatsby Charitable Foundation (GAT3361 to T.B. and T.W.M.) and Wellcome Trust (090843/F/09/Z to T.B. and T.W.M., 214333/Z/18/Z to T.W.M.). This manuscript was typset using a modified version of the HenriquesLab bioRxiv template.

## Author Contributions

S.K. and T.W.M. conceived and led the project. S.K., E.B., and T.W.M. designed experiments. D.C. and T.B. designed and built the open field recording setup. E.F.B. performed the perceptual experiments. S.K. and E.F.B. analysed the behavioural data. S.K. and D.C. performed the chronic probe implant surgeries. S.K. performed all other experiments and data analyses and designed the analysis of neural data. R.A.F. developed the “spikeGUI” analysis software. A.L.T. developed the “opendirection” analysis pipeline. D.C. and S.C.L. contributed to the development of data processing and analysis codes. T.B. and T.W.M. secured funding and resources. S.K. wrote the original draft. S.K. and T.W.M. revised and edited the manuscript with input from all the authors.

## Supplementary Tables and Figures

**Table S1.** Parameters used for automated spike sorting with KiloSort.

| Parameter | Value |
| --- | --- |
| ops.Nfilt | 640* |
| ops.whitening | full |
| ops.nSkipCov | 1 |
| ops.whiteningRange | 32 |
| ops.criterionNoiseChannels | 0.1 |
| ops.Nrank | 3 |
| ops.nfullpasses | 6 |
| ops.maxFR | 20000 |
| ops.fshigh | 300 |
| ops.ntbuff | 64 |
| ops.scaleproc | 200 |
| ops.Th | [4 10 10] |
| ops.lam | [5 20 20] |
| ops.nannealpasses | 4 |
| ops.momentum | 1./[20 400] |
| ops.shuffle_clusters | 1 |
| ops.mergeT | 0.1 |
| ops.splitT | 0.1 |
| ops.initialize | fromData |
| ops.spkTh | -4 |
| ops.loc_range | [3 1] |
| ops.long_range | [30 6] |
| ops.maskMaxChannels | 5 |
| ops.crit | -.65 |
| ops.nFiltMax | 10000 |

\*Neuronexus Poly2 = 64

**Table S2.**
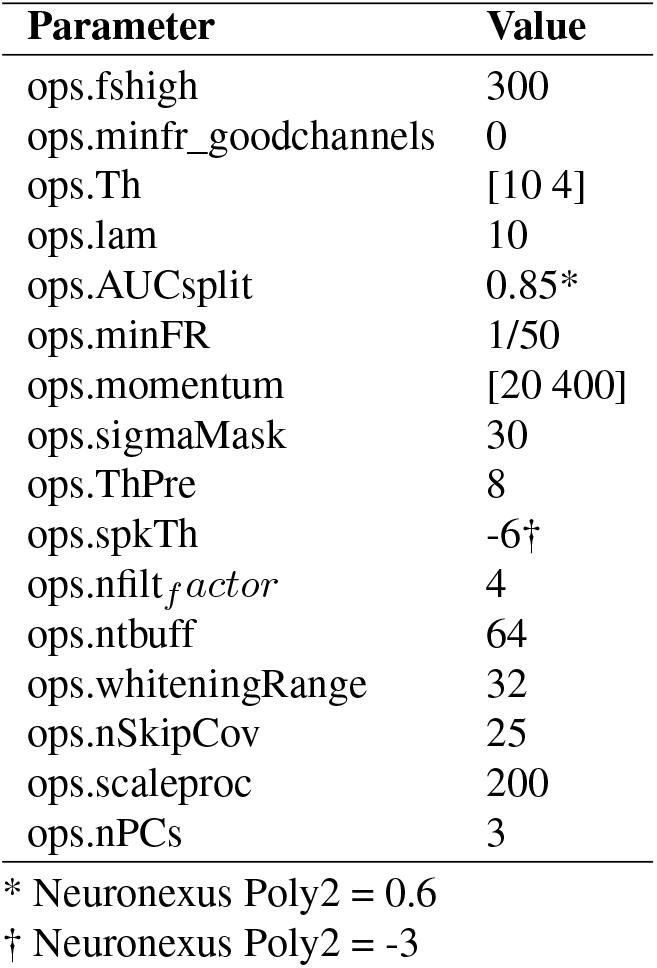
Parameters used for automated spike sorting with KiloSort2.

| Parameter | Value |
| --- | --- |
| ops.fshigh | 300 |
| ops.minfr_goodchannels | 0 |
| ops.Th | [10 4] |
| ops.lam | 10 |
| ops.AUCsplit | 0.85* |
| ops.minFR | 1/50 |
| ops.momentum | [20 400] |
| ops.sigmaMask | 30 |
| ops.ThPre | 8 |
| ops.spkTh | -6† |
| ops.nfilt<sub>factor</sub> | 4 |
| ops.ntbuff | 64 |
| ops.whiteningRange | 32 |
| ops.nSkipCov | 25 |
| ops.scaleproc | 200 |
| ops.nPCs | 3 |

\* Neuronexus Poly2 = 0.6
† Neuronexus Poly2 = -3

**Fig. S1.**
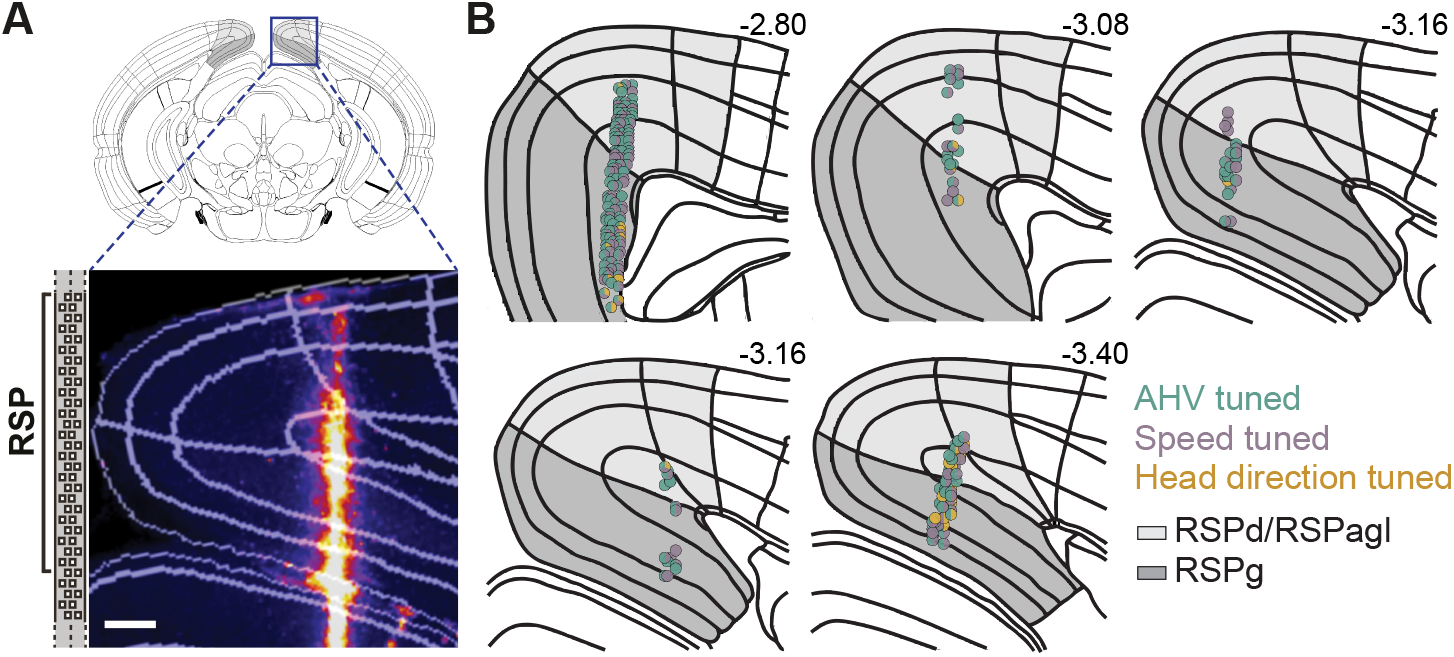
Anatomical location of chronic recordings in the RSP. **(A)** Top, schematic of a coronal brain section with RSP highlighted in grey, Bottom, inset shows a coronal 2P image of the Neuropixels probe track marked with DiI. Scale bar = 200 µm. **(B)** Schematic showing chronic recording sites in all 5 mice. Circles indicate the approximate location of cells tuned to AHV, speed, and heading direction within the dysgranular/agranular (light grey) and granular (dark grey) regions of the RSP. Numbers are distances from Bregma (in mm). All schematics and boundary outlines are based on the Allen Mouse Brain Atlas.

**Fig. S2.**
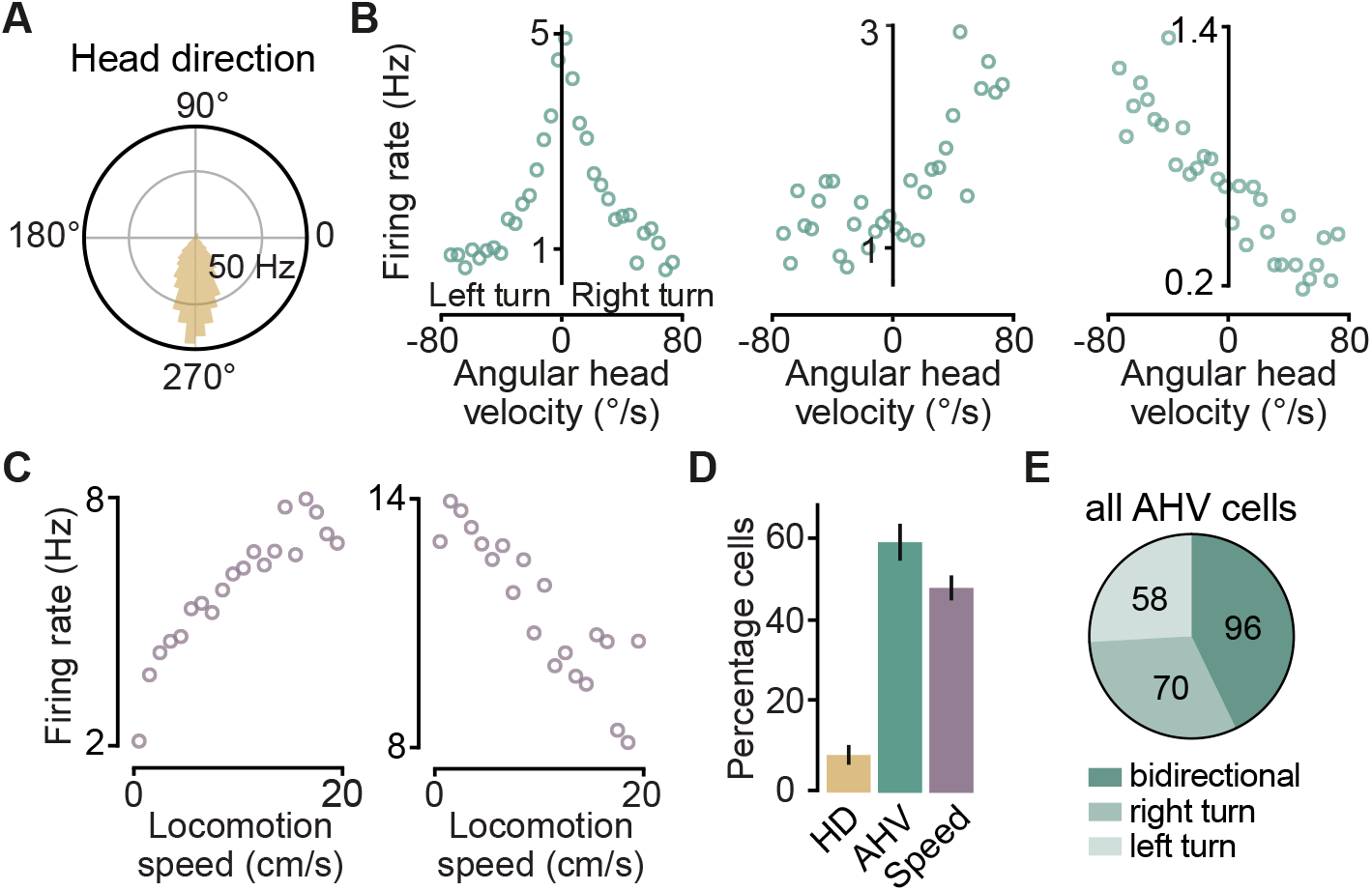
Angular head velocity, head direction, and linear speed tuning in the RSP. **(A)** An example HD cell in the RSP. Visual landmark was located at 180°. **(B)** Tuning plots for three types of AHV cells: bidirectional negative speed correlation (left), unidirectional positive correlation (centre), and opposite correlations (right). Example of a bidirectional positively correlated type is shown in Figure 1. **(C)** Tuning plots of two representative speed cells showing positive (left) and negative (right) correlations with linear locomotion speed. **(D)** Summary data (mean ± SEM, n = 5 mice, 12 recordings) showing percentage of cells that were tuned to HD, AHV, or linear speed. **(E)** Pie chart represents proportion of AHV cells (n = 224) with bidirectional and unidirectional (right or left turn) tuning.

**Fig. S3.**
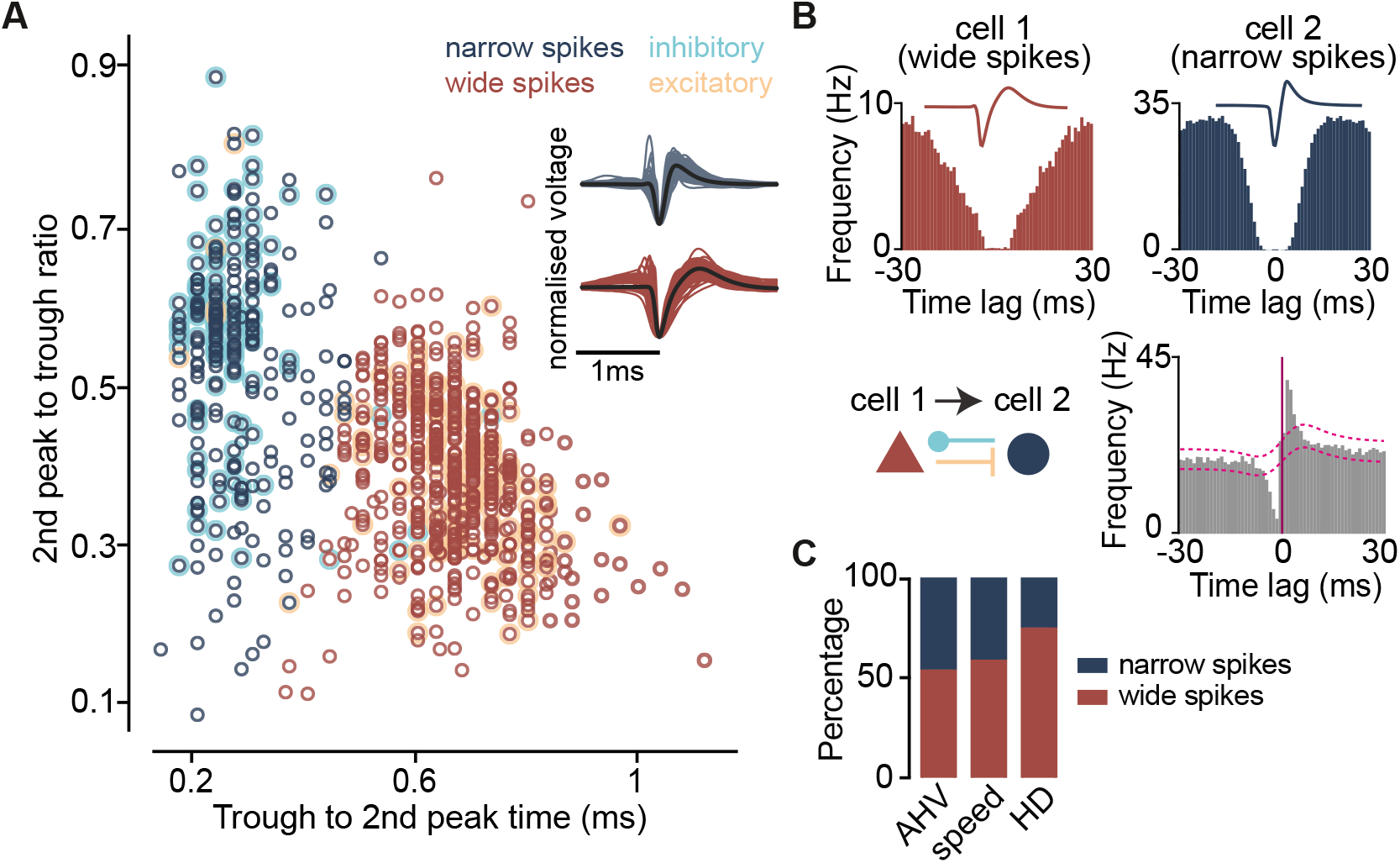
Both excitatory and inhibitory neurons in the RSP represent AHV. **(A)** K-means clustering of isolated RSP units based on spike width and peak/trough ratio. Single units were separated into two clusters with wide (putative excitatory, n = 460, 32 %, red) and narrow (putative inhibitory, n = 216, 68 %, dark blue) spikes. Cyan and orange circles mark neurons identified as inhibitory and excitatory based on cross-correlogram (CCG) analysis. **(B)**Top, average spike waveform and auto-correlogram of two example single units with wide and narrow spikes. Bottom, schematic of reciprocal monosynaptic connections (left) based on the CCG for the same pair (right). Dashed lines indicate 99.98 % confidence interval. **(C)** Percentage of putative excitatory (wide spikes) and inhibitory (narrow spike) cells among AHV (n = 224), speed (n = 182), and HD (n = 36) neurons.

**Fig. S4.**
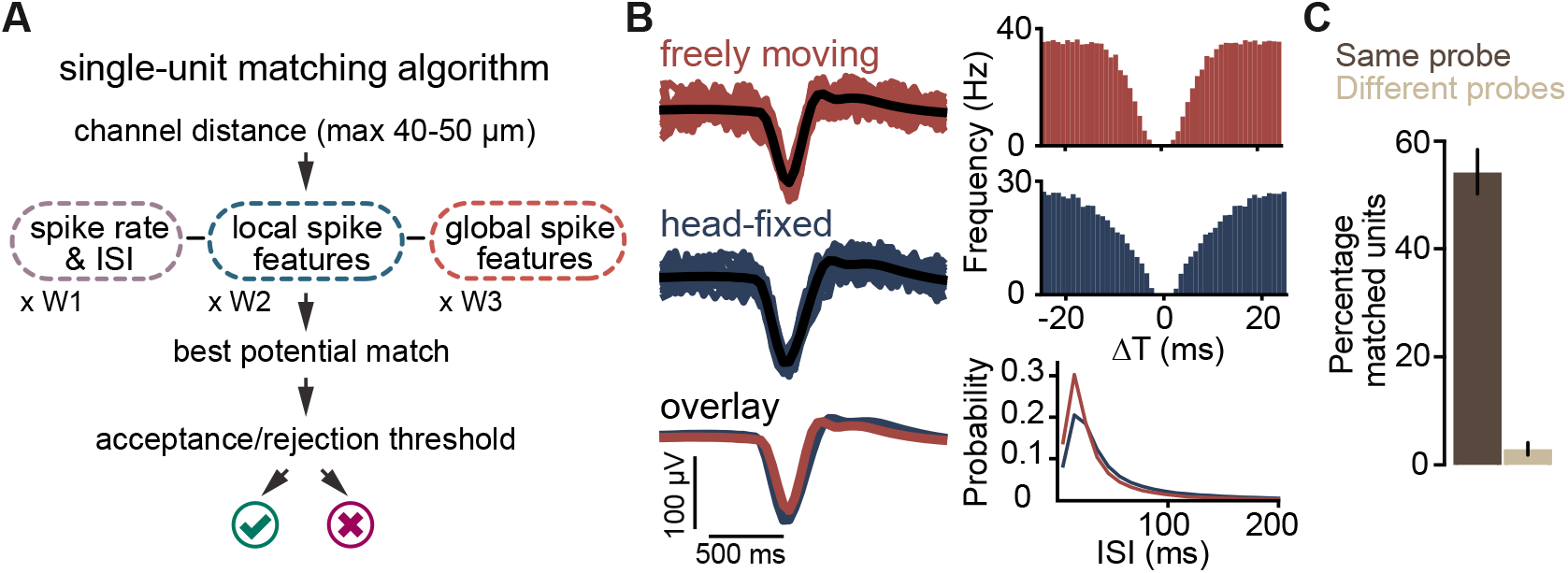
Tracking single units between freely moving and head-fixed conditions. **(A)** Schematic of the single-unit matching algorithm. Probe geometry and multiple similarity metrics for spike features, spike rate, and inter-spike interval (ISI) distribution were used to find best potential matches. **(B)** Example of a single unit tracked between head-fixed (blue) and freely moving (red) conditions. Top- and centre-left, red and blue traces are individual spikes (20 displayed). Black traces show average waveforms from all spikes. Bottom-left, average waveforms superimposed. Right, auto-correlogram (top and centre) and ISI distribution (bottom) of the tracked unit under the two conditions. **(C)** Summary data (mean ± SEM, 12 fix-free recording pairs) showing the percentage of head-fixed single units with an accepted match in the freely moving recording (same probe). Control data were generated by applying the same matching algorithm to recordings from different animals (different probes, 10 pseudo-random pairs).

**Fig. S5.**
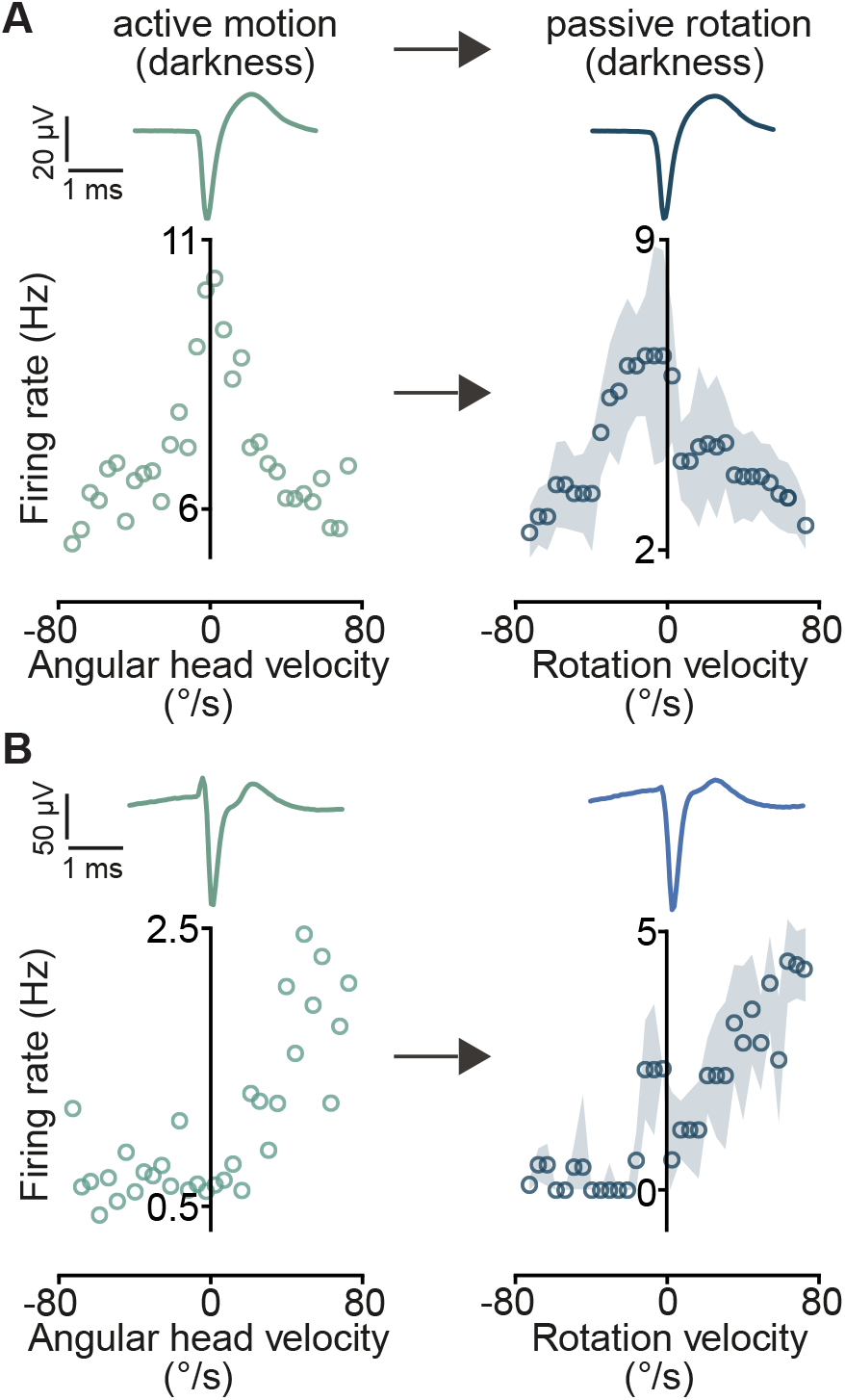
AHV neurons are similarly tuned under freely moving and head-fixed conditions. **(A)** Example of a bidirectional negatively correlated AHV cell, and **(B)** a unidirectional AHV cell recorded during both open-field exploration (left) and head-fixed passive rotation (right) in darkness. Top traces show average spike waveforms. Circles and shaded area on the right show trial-averaged firing rates (12 trials) and SEM, respectively.

**Fig. S6.**
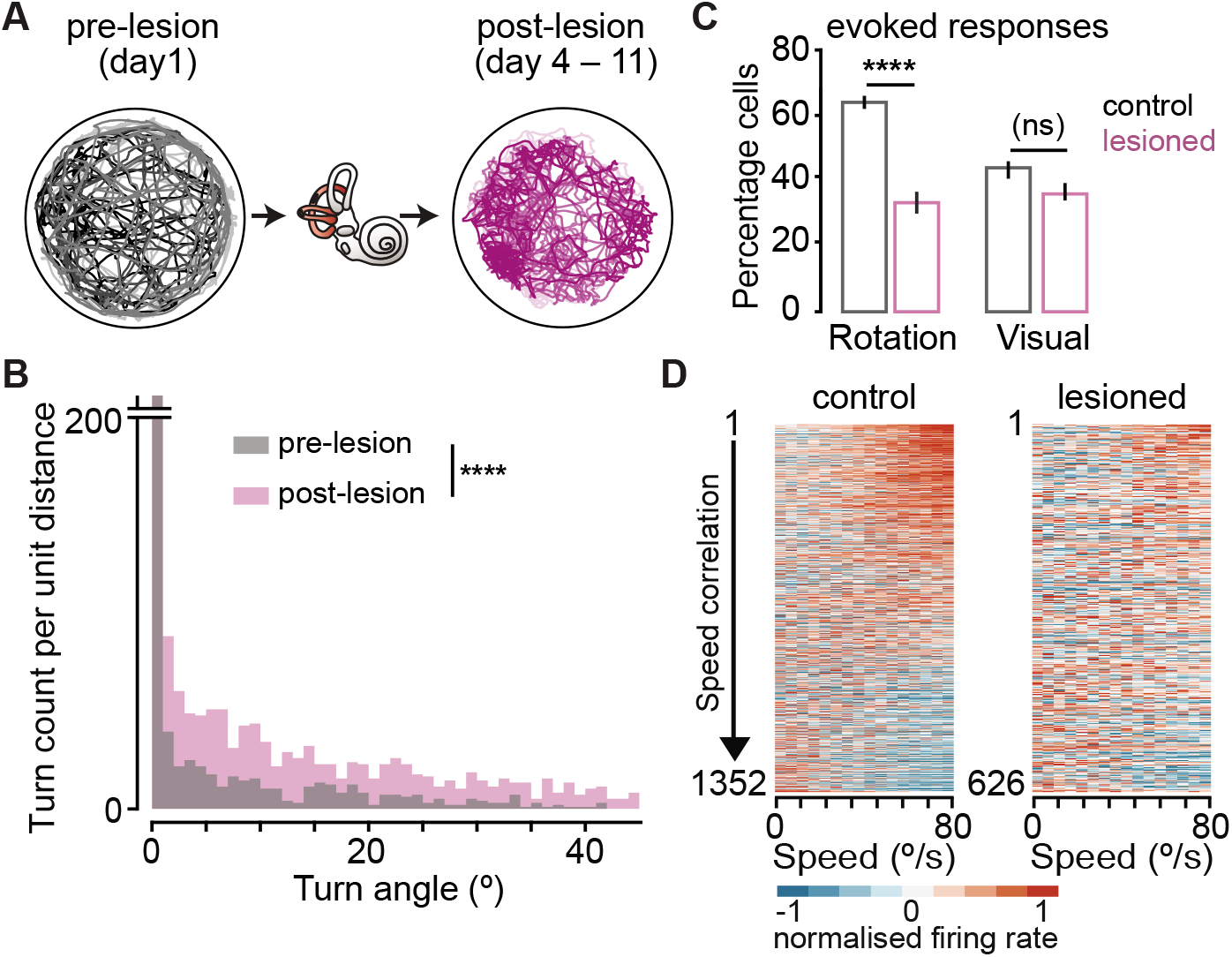
Rotation-evoked responses depend on vestibular inputs. **(A)** Overlaid open-field trajectories of 4 mice before (left) and after (right) bilateral kanamycin-induced lesions of the horizontal and posterior semi-circular canals. Lesions were confirmed by a typical increase in turning behaviour during locomotion (right and B). **(B)** Scaled histogram of turning angles, binned at 1°, for all 4 mice before (grey) and after (pink) vestibular lesions. **** P = 2e-83, Kolmogorov–Smirnov test. **(C)** Summary data (mean ± SEM; Control: n = 10 mice, 20 recordings, 676 cells; Lesioned: n = 4 mice, 7 recordings, 313 cells) showing the percentage of cells with evoked responses (significant increase or decrease in firing rate relative to baseline) to rotation in darkness or to simulated optic flow (“Visual”). **** P = 2.8e-5, (ns) P = 1, Kruskal-Wallis with Dunn’s test. **(D)** Heatmaps show baseline-subtracted, normalised average firing rate firing rate as a function of rotation speed (darkness) in controls and lesioned animals. Each pair of rows represent an individual neuron’s response to both directions and are sorted by the magnitude of speed correlations, averaged over the two directions.

**Fig. S7.**
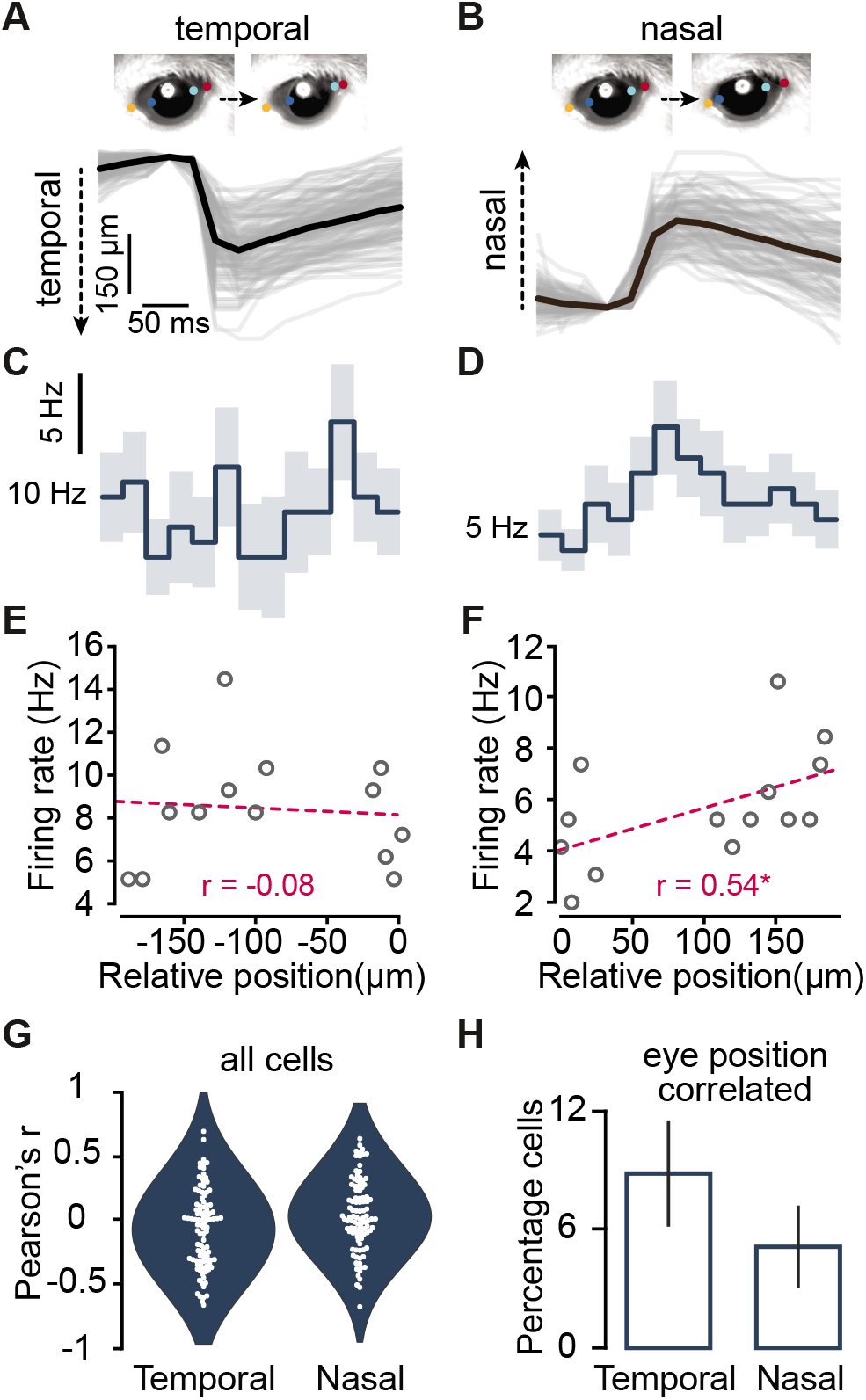
Eye movement-related activities in the RSP. **(A-B)** Top, video frames showing example temporal (A) and nasal (B) eye movements recorded at 40 fps in the dark. Coloured circles show marked positions for eye tracking using DLC. Bottom, example individual (grey), and averaged (black) eye movement events from one mouse. **(C-D)** Mean (lines) and SEM (shades) firing rate histograms of an example cell aligned with detected temporal and nasal eye-movements in A and B. Spiking rates during each eye-movement event (from 75 ms before movement onset to 250 ms after) were determined in 25 ms time bins. **(E-F)** Mean firing rate of the same cell (C-D) at each 25 ms time bin plotted against the averaged relative eye position. Dashed line shows linear fits. *Significant Pearson correlation (P = 0.048). **(G)** Population data of correlation magnitudes. Circles show Pearson’s r of correlation between firing rate and eye position for individual cells. **(H)** Summary data (mean ± SEM) showing proportion of cells with significant eye-position correlations (n = 3 mice, 101 cells, P significance threshold = 0.05).

**Fig. S8.**
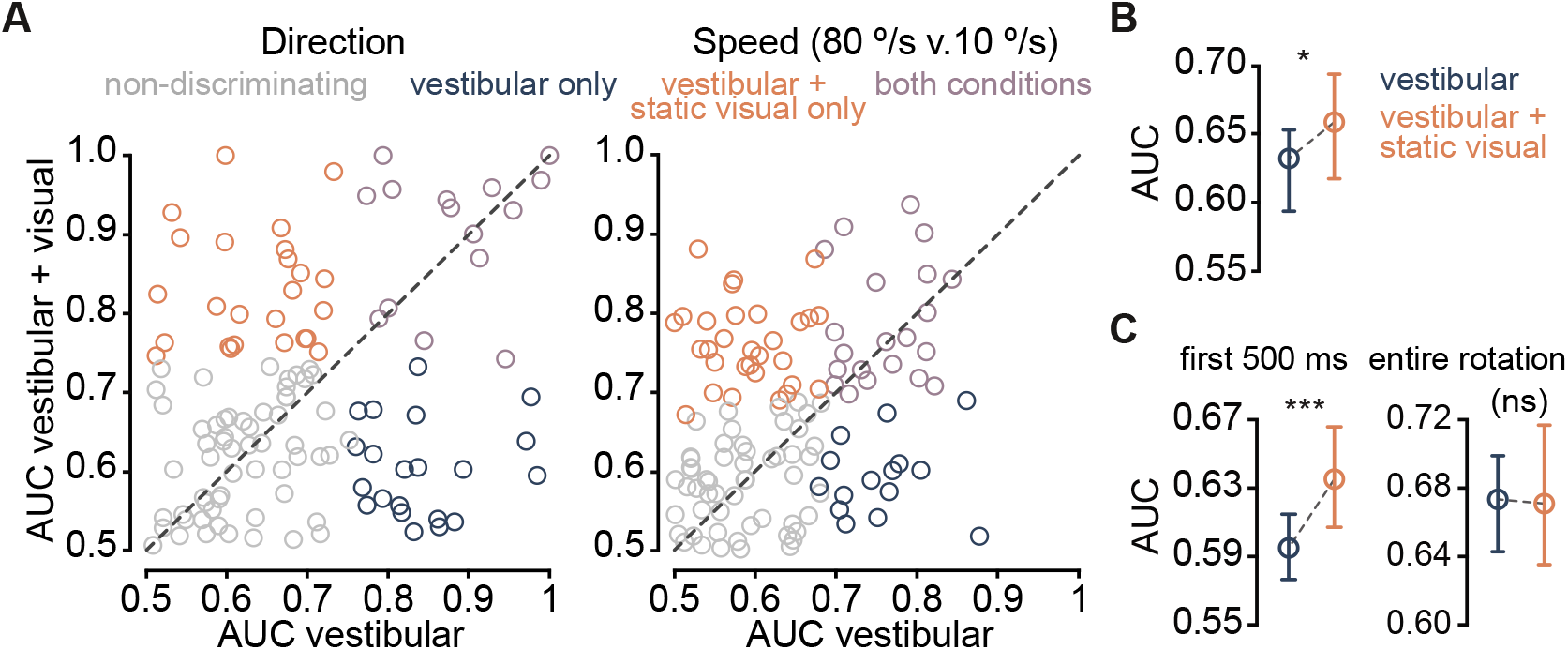
Heterogenous vestibular-visual response properties of AHV neurons. **(A)** Scatterplot of AUCs for direction (left) and speed (right) discrimination. AUCs under vestibular-visual condition (rotation with static visual stimulus) are plotted against AUCs under vestibular condition (rotation in the dark). Each circle represents one tracked AHV cell (n = 120). Direction ROC curves compared the firing rates between CW and CCW rotations. Speed ROC curves compared the firing rates between the speed bin that peaked at 10 °/s and successive speed bins (15 °/s – 80 °/s peak). Grey circles show non-discriminating cells. Blue and orange represent, respectively, cells that discriminated direction/speed exclusively in the dark (“vestibular only”), or only when visual stimuli were available (“vestibular + static visual only”). Purple indicates significant discrimination under both conditions. **(B)** Summary data (median and 95 % CI) showing percentage of cells that discriminated speed of rotation (10 °/s v. 80 °/s) under each condition. * P = 0.02, Wilcoxon signed rank test. **(C)** Summary data (median and 95 % CI) showing percentage of cells that discriminated direction of rotation under each condition, using either the first 500 ms (left), or the entire rotation window (right, 3.5 s). *** P = 3e-4, (ns) P = 0.73, Wilcoxon signed rank test.

**Fig. S9.**
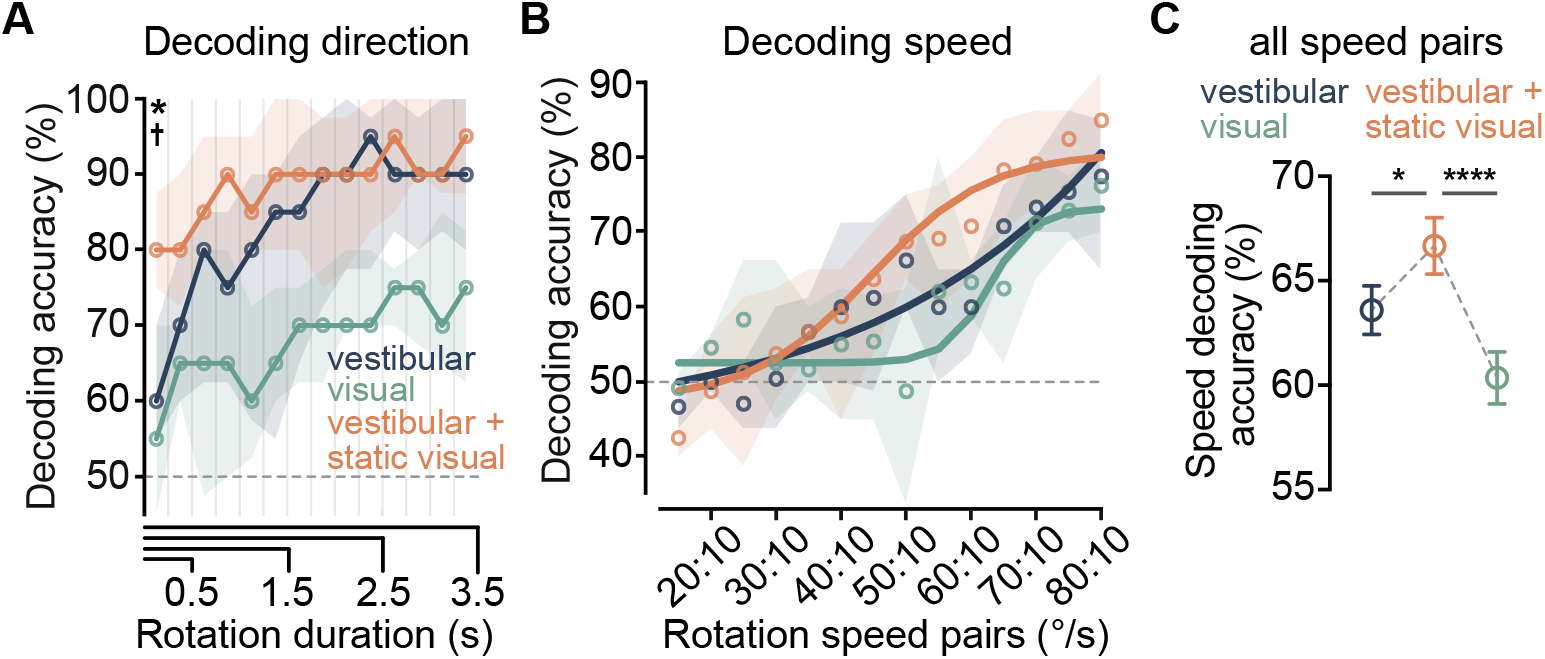
Combination of vestibular and visual cues improves decoding of angular self-motion by RSP neuronal populations. **(A)** Population decoding accuracy for direction as a function of rotation duration. Circles and shaded areas show median and IQRs. Dashed line represents theoretical chance. * P [vestibular v. vestibular + visual] = 0.04, † P [visual v. vestibular + visual] = 0.01, pairwise Wilcoxon signed rank test with Bonferonni correction, first 250 ms, n = 10 mice, 19 recordings, 10 – 84 cells per recording. **(B)** Population decoding accuracies for speed. Circles and shaded areas show median and IQRs. Lines are sigmoid fits. n = 7 mice, 12 recordings, 22 – 84 cells per recording. **(C)** Mean (± SEM) speed decoding accuracy of all speed pairs. * P = 0.01, **** P = 6.4e-5, one-way ANOVA with Holm-Sidak’s test.

